# Chromosome-scale haplotype genome assemblies for the Australian mango ‘Kensington Pride’ and a wild relative, *Mangifera laurina*, provide insights into anthracnose-resistance and volatile compound biosynthesis genes

**DOI:** 10.1101/2025.11.24.690335

**Authors:** Upendra Kumari Wijesundara, Agnelo Furtado, Ardashir Kharabian Masouleh, Natalie L Dillon, Heather E Smyth, Robert J Henry

## Abstract

Mango (*Mangifera indica*) is one of the most popular fruits cultivated in tropical and subtropical regions of the world. The availability of reference genomes helps to identify the genetic basis of important traits. Here we report assembled high-quality chromosome-level genomes for the Australian mango cultivar ‘Kensington Pride’, and *M. laurina;* a wild relative, which shows resistance to anthracnose disease. PacBio HiFi sequencing with higher genome coverage enabled the assembly of both genomes with 100% completeness. Genome sizes of ‘Kensington Pride’ and *M. laurina* were 367 Mb and 379 Mb, respectively, with all 20 chromosomes in both genomes having telomeres at both ends. K-mer analysis revealed that these genomes are highly heterozygous and significant structural variations were identified between ‘Kensington Pride’, *M. laurina,* and the recently published genome of the cultivar ‘Irwin’. Functional annotation identified key genes involved in carotenoid, anthocyanin, and terpenoid biosynthesis, responsible for fruit color and flavour in mango. Furthermore, the presence of a SNP in β-1,3-glucanase 2 gene associated with anthracnose resistance was analyzed. Whole genome duplication analysis confirmed that mangoes have undergone two polyploidization events during their evolution. Analysis revealed a conserved pattern of colinear genes, although many colinear blocks were also identified on non-homologous chromosomes.

**Practitioner Points:**

- PacBio HiFi sequencing and high coverage produced genomes for ‘Kensington Pride’ mango and *M. laurina* with 100% completeness, identifying all the telomeres in the assembled chromosomes.
- Significant structural variations were identified between ‘Kensington Pride’, *M. laurina*, and the published ‘Irwin’ genome.
- Genes linked in the biosynthesis of unique terpenoids were identified, and the structural differences in the annotated *β-1,3-glucanase 2* genes associated with anthracnose resistance provide a resource for gene expression analysis in susceptible and resistant cultivars.

## Introduction

Mango is one of the most important tropical fruits well known for its delicious taste, unique flavor, and high nutritional content. *Mangifera indica* to which all commercially growing cultivars belong is believed to have originated in North-Eastern India, the Indo-Burma region, and Bangladesh^[1]^ and then gradually spread into tropical and sub-tropical regions of the world. To date, mangoes are cultivated in more than 100 countries. In 2022, global mango production reached 44.4 million tonnes, with India accounting for 44.2% of production followed by Indonesia (9.3%), China (6.7%), Pakistan (4.7%), and Mexico (4.2%)^[2]^. Over the years, various cultivars have been selected showing wide variations in fruit quality and yield. However, there remains a continuous need for new varieties to meet evolving market demands and consumer preferences.

Breeding is the key strategy for developing new mango cultivars with high productivity, and improved fruit quality with other desired traits such as dwarfness, regular bearing habit, and biotic and abiotic stress resistance^[3,4]^. Fruit colour and flavour are major quality traits considered in mango breeding. The characteristic flavours of mango are influenced by different combinations of sugars, acids, and aroma volatile compounds, including terpenes, alcohols, esters, and lactones. Consumer preference often leans toward fruits with yellow colour skin and orange to pink, red, or purple blush. The predominant pigments that give mangoes their appealing skin and blush colours are carotenoids and anthocyanins respectively^[5]^. Breeding for dwarf varieties, tolerance to marginal soils and saline water are important in increasing tree density and minimizing resource use and costs. Furthermore, resistance to pre- and post-harvest diseases is highly desirable to improve fruit quality and increase the yield^[3]^. Although mango breeding is slow due to some of the inherent traits such as a long juvenile phase, polyembryony, and high heterozygosity, advances in genome sequencing technologies have enabled the assembly of high-quality reference genomes for parental genotypes. Developing such comprehensive genomic resources allows researchers to identify key genes associated with desirable traits and develop molecular markers to accelerate mango breeding. Marker-assisted selection helps identify individuals with desired traits, reducing the need to maintain a large breeding population over long periods^[3,6]^.

*M. indica* cv. ‘Kensington Pride’ is the most widely grown mango variety in Australia, with significant consumer acceptance due to its distinctive aroma and flavour. This unique flavour profile is primarily determined by volatile compounds, including monoterpenes (49%), esters (33%), and lactones. Among these, the volatile compound α-terpinolene has been identified as the most abundant monoterpene contributing to its unique flavour^[7]^. ‘Kensington Pride’ also possesses other favourable attributes, such as wide adaptability to agroclimatic conditions, and an attractive appearance, making it the main parental variety used in the Australian mango breeding program. However, it also has problems including irregular bearing, high vigor, and susceptibility to diseases^[8]^.

Crop wild relatives are potential sources of allelic variation that help crops overcome biotic and abiotic stresses. High-quality genomes of crop wild relatives can be used to explore genes and quantitative trait loci associated with agronomically important traits for crop improvement^[9]^. Several wild relatives of mango producing edible fruits have been identified with traits that may be useful in breeding programs. Among them, *M. laurina* exhibits resistance to anthracnose, a major pre- and post-harvest fungal disease that significantly affects mango yield. *M. laurina* is well adapted to grow in wet and humid environments and can thrive in areas where common mangoes struggle due to susceptibility to anthracnose, resulting in poor fruit set. The species identified so far in the genus *Mangifera*, including cultivated mango, are diploid (2n=40) and crosses between *M. indica* and *M. laurina* have successfully resulted in 60 hybrids^[10]^.

A recently published genome for the mango cultivar ‘Irwin’ demonstrated that PacBio HiFi data together with high genome coverage alone can produce highly contiguous reference genomes^[11]^. In this study, using HiFi sequencing together with high genome coverage, we developed high-quality genomes for the cultivar ‘Kensington Pride’ and the wild relative *M. laurina*. All the chromosomes of both genomes were assembled with telomeric repeats at both ends, indicating assembly of full-length chromosomes. The Comparison of the ‘Kensington Pride’ and *M. laurina* genomes with the recently published genome of the cultivar, ‘Irwin’^[11]^ identified significant structural variations among the three genomes. Functional annotation identified key genes associated with the biosynthesis of aroma volatile compounds and disease resistance. Therefore, the genomes assembled in this study provide a valuable resource for understanding the genetic basis of important traits in mango breeding.

## Results

### Genome sequencing and assembly

The total yields of HiFi data for ‘Kensington Pride’ and *M. laurina* were 79.46 Gb (217x coverage) and 76.07 Gb (201x coverage), respectively (Table S1). For each species, the HiFiasm tool generated a collapsed assembly and two haplotypes. The ‘Kensington Pride’ collapsed assembly, haplotype 1 (hap1) and haplotype 2 (hap2) consisted of a total of 4387, 4499, and 1380 contigs, respectively, while the *M. laurina* collapsed, hap1 and hap2 assemblies were composed of 3899, 4159 and 1301 contigs. We recently published a high-quality reference genome for *M. indica* cv. ‘Irwin’^[11]^, which had assembly completeness of 100% assessed by Benchmarking Universal Single-Copy Orthologs (BUSCO) analysis. The collapsed genomes assembled here for ‘Kensington Pride’ and *M. laurina* also showed 100% completeness whereas the haplotype assemblies of both species showed more than 98% completeness (Table 1). Furthermore, the collapsed assemblies for ‘Kensington Pride’ and *M. laurina* had contig N50s of 15.05 Mb and 15.93 Mb, respectively, showing even higher assembly contiguities than the Irwin genome. In addition, K-mer analysis revealed that the *M. laurina* assembly showed the highest heterozygosity (2.22%), while ‘Kensington Pride’ showed higher heterozygosity (1.77%) compared to ‘Irwin’ (1.24%) (Figure S1).

**Table 1.** Comparison of ‘Irwin’, ‘Kensington Pride’, and *M. laurina* genome assembly and annotation.

| Statistic | 'Irwin'* |  |  | 'Kensington Pride' |  |  | *M. laurina* |  |  |
| --- | --- | --- | --- | --- | --- | --- | --- | --- | --- |
|  | Collapsed | Hap 1 | Hap 2 | Collapsed | Hap 1 | Hap 2 | Collapsed | Hap 1 | Hap 2 |
| **Assembly** |  |  |  |  |  |  |  |  |  |
| Total number of contigs | 4,642 | 4,711 | 1,515 | 4,387 | 4,499 | 1,380 | 3,899 | 4,159 | 1,301 |
| Total length (bp) | 556,539,885 | 543,231,508 | 432,295,890 | 549,901,289 | 539,878,467 | 435,195,159 | 558,942,218 | 552,234,473 | 454,644,896 |
| GC % | 36.50 | 36.52 | 34.77 | 36.64 | 36.58 | 35.09 | 36.26 | 36.13 | 34.58 |
| Genome size (20 chromosomes only) | 365 | 354 | 355 | 367 | 361 | 358 | 379 | 367 | 381 |
| Contig N50 (Mb) | 14.98 | 13.21 | 15.45 | 15.05 | 13.76 | 13.75 | 15.93 | 14.75 | 15.82 |
| Contig L50 | 14 | 16 | 12 | 15 | 15 | 12 | 14 | 14 | 12 |
| Number of telomeres | 40 | 40 | 39 | 40 | 39 | 40 | 40 | 40 | 40 |
| Complete BUSCO (%) (Viridiplantae, N=425) | 100 | 98.1 | 99.0 | 100 | 100 | 100 | 100 | 99.8 | 100 |
| **Annotation** |  |  |  |  |  |  |  |  |  |
| Annotation completeness (N=425, %) | 99.6 | 97.6 | 98.6 | 99.1 | 98.8 | 99.6 | 99.5 | 99.3 | 99.5 |
| Number of protein-coding genes | 35,220 | 34,659 | 33,230 | 34,361 | 34,651 | 34,229 | 34,278 | 34,717 | 35,216 |
| Number of CDS sequences | 42,973 | 42,268 | 40,947 | 41,779 | 41,916 | 41,216 | 41,488 | 41,912 | 42,584 |
| Number of proteins | 42,973 | 42,268 | 40,947 | 41,779 | 41,916 | 41,216 | 41,488 | 41,912 | 42,584 |
| Total gene length (bp) | 110,918,018 | 107,399,304 | 108,372,140 | 110,701,932 | 109,430,362 | 107,064,408 | 111,517,258 | 109,472,501 | 111,169,860 |
| Total exon length (bp) | 52,229,406 | 51,040,974 | 50,850,835 | 51,034,566 | 51,045,231 | 49,985,022 | 50,340,320 | 50,175,744 | 50,902,422 |
| mean gene length (bp) | 3,149 | 3,098 | 3261 | 3,221 | 3,158 | 3,127 | 3,253 | 3,153 | 3,156 |
| mean exon length (bp) | 200 | 200 | 199 | 199 | 202 | 202 | 200 | 201 | 201 |
| Longest gene (bp) | 59,081 | 59,081 | 48,708 | 61,325 | 61,325 | 62,715 | 61,080 | 59,699 | 60,241 |
| Shortest gene (bp) | 201 | 201 | 228 | 201 | 201 | 201 | 219 | 201 | 201 |

The published ‘Irwin’ genome was used as a reference to orient and assign contigs of ‘Kensington Pride’ and *M. laurina* assemblies into chromosomes (Figure S2a,d). According to dot plots, 16 chromosomes of the ‘Kensington Pride’ collapsed assembly were each represented by a single contig, where all the contigs had telomeres at both ends, indicating all 16 as complete chromosomes. Chromosomes 6, 8, and 11 consisted of two contigs, and chromosome 7 consisted of three contigs. Most of the contigs in chromosomes 6, 7,8, and 11 had rRNA repeats at the ends, which required joining to get a complete chromosome. The contigs in chromosome 6 had telomeric repeats at one end and gene sequences at the other end. However, contigs in chromosomes 8 and 11 had 28S rRNA repeats at ends to be linked, and telomeric repeats at the other ends. Furthermore, two of the three contigs in chromosome 7 had telomeric repeats at one end and 5S rRNA repeats at the ends required joining, while the middle contig had repetitive sequences at both ends (Tables S2 and S3). Therefore, contigs in chromosomes 6, 7, 8, and 11 were joined by 100 Ns to generate complete pseudomolecules since they had either the same repetitive sequence at the ends that required joining and/or were aligned with the same chromosome (Figure 1e). Once the contigs were joined, the ‘Kensington Pride’ collapsed genome (367 Mb) consisted of 25 contigs and all 20 chromosomes had telomeres at both ends (Figure 1e, Table S3). BUSCO analysis of both the entire assembly with all the contigs and the assembled 20 chromosomes revealed identical and highest assembly completeness (100%). In addition, other than the contigs assembled into 20 chromosomes, the remaining 4,362 contigs in ‘Kensington Pride’ were relatively very small (15 kb - 1.1 Mb), and showed high sequence similarity to the chloroplast, mitochondrial genomes, and to the nuclear rRNA genes (Figure S2g-i). Therefore, the Kensington Pride genome consisted only of 20 chromosomes. In *M. laurina*, 16 chromosomes of the collapsed assembly were each represented by a single contig, where all the contigs had telomeres at both ends. The other four chromosomes were represented each by two contigs. Contigs of chromosomes 8, 11, and 19 had rRNA repeats at one end and telomeric repeats at the other end. In chromosome 7 only, both contigs had telomeres at one end, where one contig had 5S rRNA repeats at the other end and the other contig had a gene sequence (Tables S2 and S4). When 100 Ns joined contigs in chromosomes 7, 8, 11, and 19, all the pseudomolecules had telomeric sequences at both ends, and the genome (379 Mb) consisted of 24 contigs (Figure 1k, Table S4). Similar to ‘Kensington Pride’, both the whole contig assembly and the contigs assembled into 20 chromosomes of *M. laurina* showed 100% assembly completeness based on BUSCO analysis. In addition, except the ones assembled into 20 chromosomes, 3,875 remaining contigs in *M. laurina* were also relatively small, ranging from 16 to 796 kb. Most of these contigs showed high sequence similarity to chloroplast and mitochondrial genomes, as well as to nuclear rRNA gene sequences, a pattern consistent with the results observed for ‘Kensington Pride’ (Figure S2j-l). Therefore, final assembly of *M. laurina* consisted of 20 chromosomes only. The two haplotype assemblies of ‘Kensington Pride’ and *M. laurina* genomes were aligned with the respective collapsed genomes, and contigs belonging to the same chromosome were linked to develop complete pseudomolecules (Figure S2b,c,e,f). Though the haplotype assemblies of both ‘Kensington Pride’ and *M. laurina* were less contiguous requiring 27-35 contigs, 13-15 chromosomes were represented each by a single contig. Furthermore, except hap 1 of ‘Kensington Pride’ which had 39 telomeres, the other haplotype of Kensington Pride and each of the two haplotypes of *M. laurina* had all 40 telomeres in their genomes (Table S3,S4).

**Figure 1:**
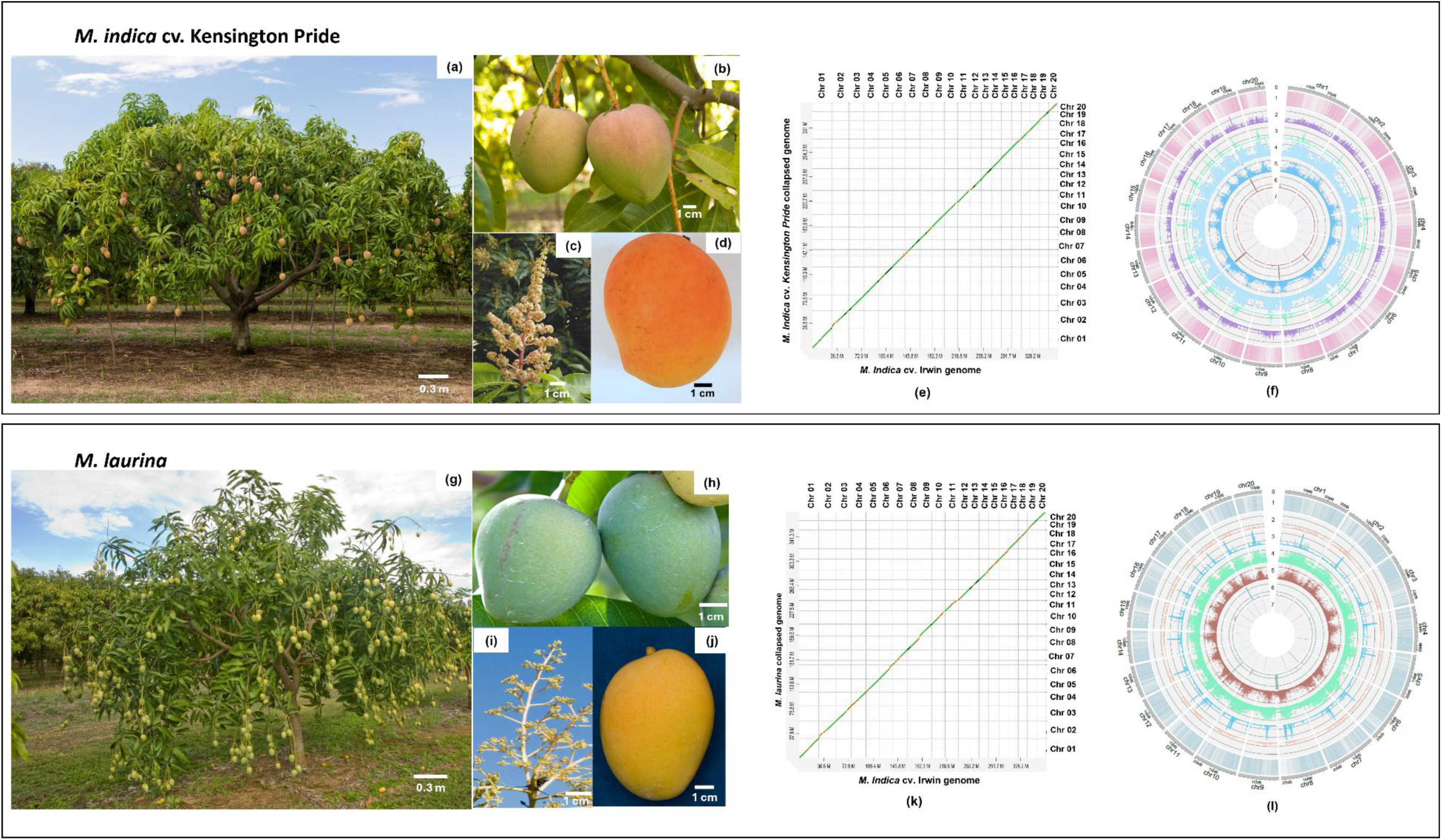
Overview of ‘Kensington Pride’ and *M. laurina* plants and genomes. (a/g): plant, (b/h): unripe fruits, (c/i): flowers, and (d/j): ripe matured fruit, (e/k): Alignment between ‘Irwin’ collapsed genome versus ‘Kensington Pride’/ *M. laurina* collapsed genome, (f/i): circos plot for ‘Kensington Pride’/ *M. laurina* collapsed genome. In the circus plots, each track with numbers indicates follows: (0) 20 pseudochromosomes (Mb), (1) predicted genes (2) Regions of DNA TE elements; (3) Regions of LINEs; (4) LTR Copia elements; (5) Regions of LTR Gypsy elements; (6) Regions of ribosomal RNA, tRNA, and snRNA repetitive regions; and (7) Telomeric repeats.

### Repetitive element identification, gene prediction, and functional annotation

For both species, only the 20 assembled chromosomes were considered for the annotations as assembly completeness was identical for the entire assembly and the 20 chromosomes. The sizes of the chromosomes and the number of genes in collapsed, hap1, and hap2 genomes are included in Table 2. Repetitive sequence analysis revealed that the ‘Kensington Pride’ collapsed genome had a higher repetitive content (49.4 %) compared to the ‘Irwin’ collapsed genome (48.7%)^[11]^. However, the *M. laurina* collapsed genome had the highest repetitive content (51.1%) when compared to the two *M. indica* cultivars. A large portion of the genomes were covered by interspersed repeats (‘Irwin’: 46.3%, ‘Kensington Pride’: 46.7%, *M. laurina*: 48.1%). Unclassified repeats were the predominant repeats among different types of repetitive sequences, while the most prevalent classified repeats in all three genomes were long terminal repeat (LTR) elements (Figure 1, Table S5, S6). A total of 70.45 Gb (192x coverage) and 119 Gb (313x coverage) RNA sequence reads of ‘Kensington Pride’ and *M. laurina* were used for annotating protein-coding genes. Gene prediction in Braker resulted in 34,361 and 34,278 genes in the ‘Kensington Pride’ and *M. laurina* collapsed genomes (in 20 chromosomes) with 41,779 and 41,488 protein sequences, respectively. The total number of genes of the collapsed genomes, hap1 and hap2, (of the 20 chromosomes) are included in Table 1. The completeness of the annotated genes was also high for all the genomes (Table 1). During functional annotation, 94.5% and 94.4% of the genes in ‘Kensington Pride’ and *M. laurina* collapsed genomes had blast hits while it was 94.1-94.3% and 93.7-93.8% for the haplotypes, respectively (Figure S3). Furthermore, the majority of the genes that didn’t have any blast hit had coding potential (Figure S4). In total, 75.5% and 75.4% of the protein-coding genes in the ‘Kensington Pride’ and *M. laurina* collapsed genomes were functionally annotated, respectively, whereas in haplotypes, 75.5 - 75.6%, and 74.7-74.8% genes were annotated (Figure S3).

**Table 2.**
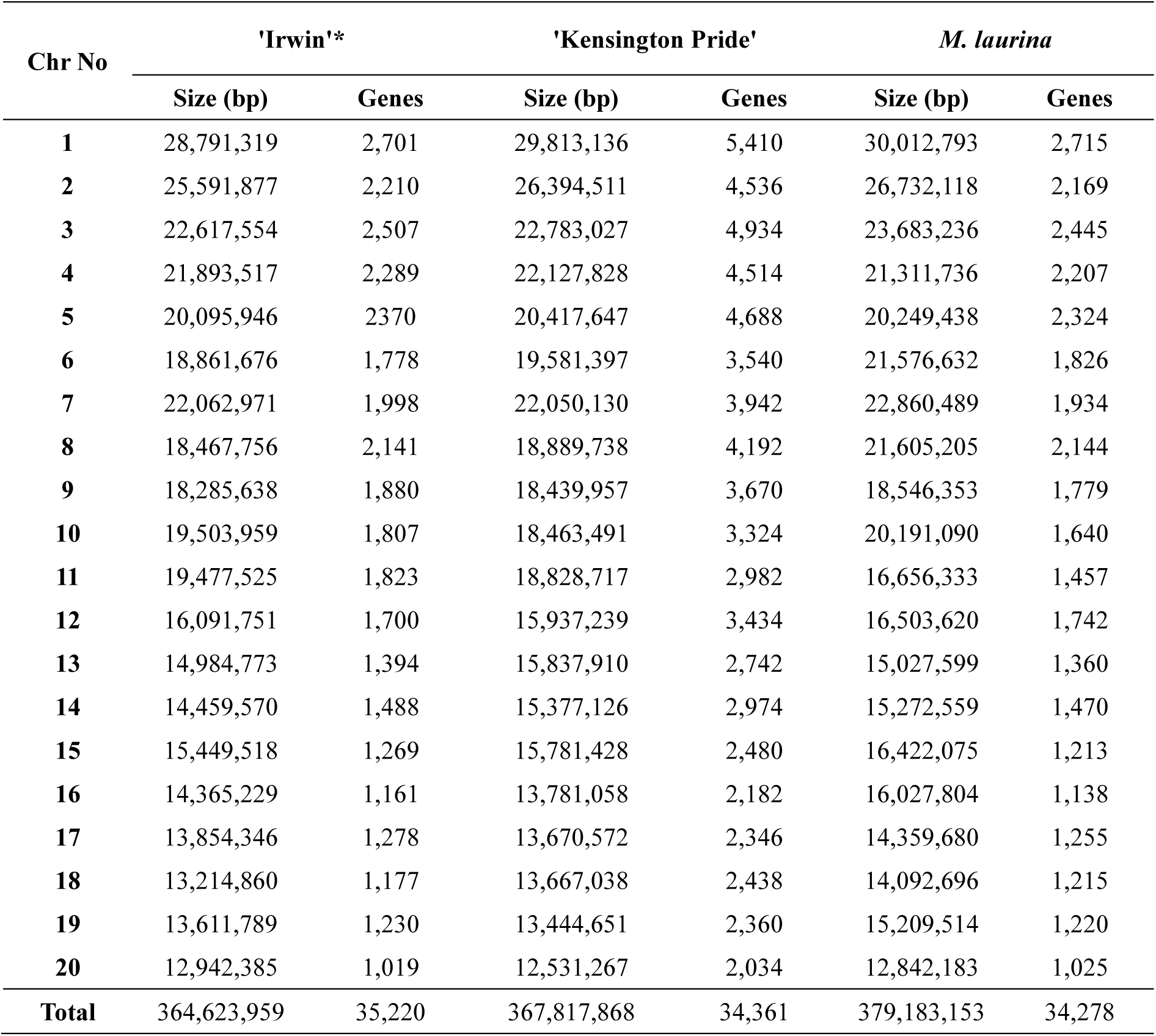
Sizes of the chromosomes and the number of genes in ‘Kensington Pride’, *M. laurina* genomes and previously published ‘Irwin’ genome.

### Genome comparison for structural variations

’Kensington Pride’, ‘Irwin’ and *M. laurina* genomes were analyzed using Syri^[12]^ to identify structural variations (Figure 2). Comparison of ‘Kensington Pride’ and ‘Irwin’ genomes identified 312.7-312.8 Mb of syntenic regions and 2,846 translocations. Furthermore, a total of 9,220 and 7,868 duplications were detected in ‘Irwin’ and ‘Kensington Pride’ genomes, respectively, and ‘Kensington Pride’ genome had 85 inversions, 1,134 insertions, and 1,184 deletions compared to ‘Irwin’ (Table S7). When the *M. laurina* genome was aligned with the ‘Irwin’ genome, less syntenic regions (295.4-295.8 Mb) and a higher number of translocations (4,710) were identified compared to ‘Irwin’ vs. ‘Kensington Pride’. Similarly, compared to ‘Irwin’ vs. ‘Kensington Pride’ genome assessment, the syntenic region (296.3 Mb) was low and the number of translocations (4,590) was higher between the *M. laurina* and ‘Kensington Pride’ genomes. Furthermore, other structural variations including inversions, duplications, insertions and deletions were also higher in *M. laurina* when compared with the ‘Irwin’ and ‘Kensington Pride’ genomes (Table S7). Interestingly, there were inversions unique to *M. laurina* in 14 chromosomes which were not present in either of the two *M. indica* cultivars (Figure S5). Chromosomes 7 and 13 had two relatively large inversions (1 Mb and 0.6 Mb respectively) while other chromosomes had inversions ranging between 1-500 kb. Comparing haplotypes of ‘Kensington Pride’ and *M. laurina* identified high structural variations (Table S8, Figure S5).

**Figure 2:**
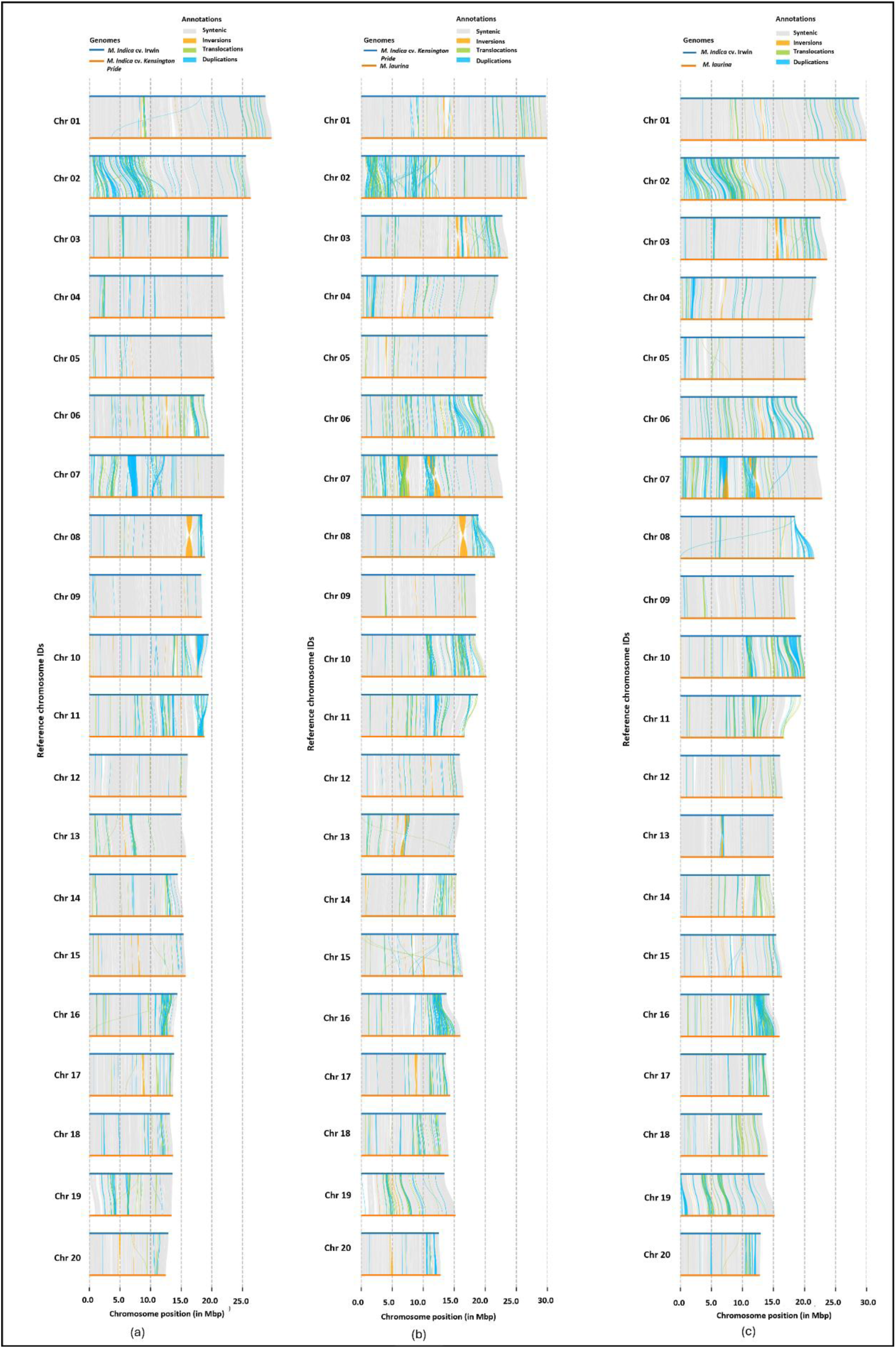
Chromosome-wise structural variations between mango collapsed genomes (a) ‘Irwin’ versus ‘Kensington Pride’ (b) ‘Kensington Pride’ versus *M. laurina and* (c) ‘Irwin’ versus *M. laurina*.

### Colinear gene analysis, duplicated gene classification, and whole genome duplication (WGD) events in mango

Collinear interactions can offer valuable insights into the evolutionary history of a genome, and it is helpful to detect evidence for WGD events and complex chromosomal rearrangements. When pair-wise collinear relationships were analyzed among the three genomes, we identified that many genes and their order were conserved between the two corresponding chromosomes. However, we also could see colinear blocks between different chromosomes detecting chromosomal rearrangements (duplications and translocations) (Figure 3a). The number of collinear genes shared between ‘Irwin’ and ‘Kensington Pride’ was 32,432 (46.6%). Similarly, ‘Irwin’ and *M. laurina* shared 33,408 (48.1%) colinear genes with 30,764 (44.8%) shared between ‘Kensington Pride’ and *M. laurina* revealing a slightly higher number of colinear genes shared between ‘Irwin’ and *M. laurina* than between the two *M. indica* cultivars. Other than the colinear blocks detected between the same chromosome, many colinear blocks were identified between chromosomes 11 and 19, 13 and 17, and 16 and 1 of all genome comparisons. The degree of collinearity in chromosome 2 was relatively low in any pair of genomes and colinear genes in chromosome 2 were rearranged with chromosomes 7 and 9. Gene duplications accounted for a significant fraction of the colinear gene rearrangements, whereas translocations accounted for the rest (Figure 3a).

**Figure 3.**
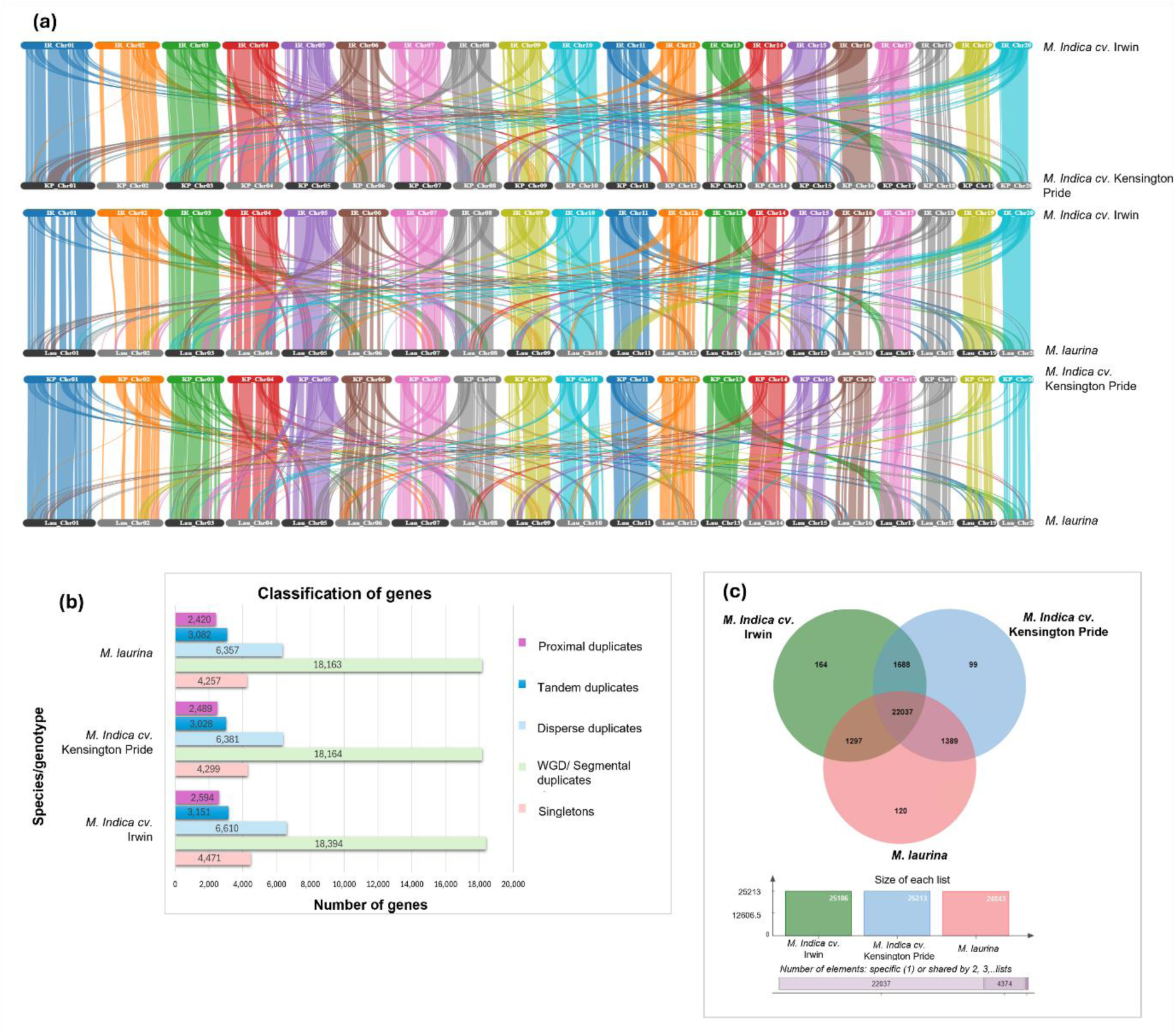
Gene collinearity, gene duplication events and unique gene cluster analysis. (a) Colinear genes between *M. indica* cv. ‘Irwin’ and ‘Kensington Pride’, ‘Irwin’ and *M. laurina*, and ‘Kensington Pride’ and *M. laurina* (b) four different mechanisms for the origin of gene duplication events in genomes. The highest number of genes originated due to whole genome duplication/segmental duplication events in all three genomes (c) Venn diagram displaying the shared and unique gene clusters among ‘Irwin’, ‘Kensington Pride’ and *M. laurina* genomes.

Based on the copy number of genes and their distribution across the genomes, all the genes were classified into singletons, dispersed duplicates, proximal duplicates, tandem duplicates, and segmental/WGD duplicates. The majority of the duplicated genes (18,163-18,394) were classified as segmental/WGD duplicates (∼52%). In each genome, dispersed duplicates (∼18%) were the second predominant type of duplicates which was followed by tandem duplicates (∼8%), and proximal duplicates (∼7%) (Figure 3b). The remaining genes present as single copies were classified into singletons representing nearly 12% of the total number of genes.

During evolutionary history, angiosperms have experienced one or more polyploidizations, and in mango, two WGD events have been identified^[13]^. Therefore, we used wgdi 0.6.5^[14]^ to estimate WGD in three genomes using median synonymous substitutions per site (Ks). The analysis of the median Ks for paralogous gene pairs in ‘Irwin’, ‘Kensington Pride’, and *M. laurina* revealed two distinct peaks, corresponding at Ks values of approximately 0.3 and 1.5 (Figure S6). This confirmed the occurrence of two WGD events in mango, as previously identified^[13]^.

### Conserved and unique gene family analysis

Analysis of unique and conserved gene families among ‘Irwin’, ‘Kensington Pride’ and *M. laurina* identified 26,794 orthogroups and 4,193 singletons (unique genes in the genomes that were not assigned to orthogroups) (Figure 3c). A total of 22,037 gene families were shared among the three genomes including 86,568 genes (’Irwin’: 28,676, ‘Kensington Pride’: 28,877, *M. laurina*: 29,015). ‘Irwin’ and ‘Kensington Pride’ shared 1,688 gene families while *M. laurina* shared 1,297 and 1,389 gene families with ‘Irwin’ and ‘Kensington Pride’, respectively. In ‘Irwin’, 2,594 unique genes were identified including 807 genes classified under 164 orthogroups and 1,787 singletons. While 1,508 unique genes (381 genes in 99 orthogroups and 1,127 singletons) were detected in ‘Kensington Pride’, 1,752 unique genes (453 genes in 120 orthogroups and 1,279 as singletons) were identified in *M. laurina*. The highest number of unique genes found in all three genomes encoded proteins that are components of the intracellular anatomical structure. Furthermore, a higher number of unique genes were found to be enriched in biological processes such as organic substances, primary and cellular metabolic processes, biosynthetic processes, and stress response. In addition, a significant number of unique genes were involved in molecular functions such as organic cyclic compound binding, small molecule binding, protein binding, hydrolase activity and transferase activity (Figure S7).

KEGG pathway analysis revealed that unique genes of both *M. indica* genotypes and *M. laurina* were mainly associated with purine and thiamine metabolism. Compared to *M. laurina* and ‘Kensington Pride’, ‘Irwin’ had a higher number of unique genes enriched in purine and thiamine metabolism (80 and 75 genes, respectively). Other than these two pathways, the unique genes in ‘Irwin’ also included those involved in plant-pathogen interaction, phenylpropanoid biosynthesis and co-factor biosynthesis. A significant number of unique genes in ‘Kensington Pride’ were enriched in anther and pollen development, response to drought, and tryptophan metabolism whereas in *M. laurina*, a relatively higher number of unique genes were engaged in pathways such as response to drought, plant-pathogen interactions, tryptophan metabolism, sesquiterpene and triterpenoid biosynthesis (Table S9).

Important genes in *M. indica* cv. ‘**Irwin’, ‘Kensington Pride**’ and *M. laurina*

### Anthracnose resistant gene

A SNP (G to A) within the β-1,3-glucanase 2 (*β-1,3-GLU2*) gene which substitutes the amino acid, isoleucine with valine in the encoded protein has been identified to enhance the defence response of the gene against *Colletotrichum gloeosporioides*, the causative fungal organism of anthracnose disease in mango^[15]^. We identified *β-1,3-GLU2* gene copies in all three genomes by searching the gene region relevant to the resistant SNP^[15]^. Two copies of the *β-1,3-GLU2* gene were identified in chr5 and chr9 of disease-resistant *M. laurina*, but only one copy of the gene (g19204 in chr 9) had the SNP responsible for disease resistance whereas the other copy (g9716 in chr5) had the SNP for the disease susceptibility. The two genes also showed structural differences in the encoded proteins. Gene g19204 encoded one protein with seven exons whereas gene g9716 encoded two protein sequences, which had three and six exons, respectively. ‘Kensington Pride’ and ‘Irwin’ exhibit disease susceptibility, but both these genomes had two copies of the *β-1,3-GLU2* gene in chr 9 and chr 15 with the disease-resistant SNP and two copies of the gene in chr 4 and chr 5 with the disease-susceptible SNP (Figure S8). In ‘Kensington Pride’, both genes with the resistant SNP (g19358 and g28189) encoded only one protein with four exons. However, each gene (g7731 and g9897) with the disease-susceptible SNP encoded two proteins where the shorter proteins had 3-4 exons and the longer proteins had 6-7 exons. Furthermore, in ‘Irwin’, genes with the disease-resistant SNP (g19510 and g28848) and the susceptible SNP (g7667 and g9892) each encoded a single protein where g19510 and g28848 had eight and five exons and g7667 and g9892 genes had four and six exons, respectively (Table S10). Therefore, although the three genotypes show different responses against anthracnose disease, all three had *β-1,3-GLU2* genes with the resistant and susceptible SNP which encoded structurally diverse proteins.

### Fruit peel and flesh coloration-related genes

Mango fruit peel colour varies from green, yellow, and orange to red. ‘Irwin’ is a red fruit cultivar, and anthocyanins are the pigments responsible for red peel. A total of 16 enzymes are involved in anthocyanin biosynthesis in mango (Figure S9). In ‘Irwin’, a total of 127 structural genes encoding phenylalanine deaminase, 4-coumarate CoA ligase, trans-cinnamate 4-monooxygenase, chalcone synthase, flavanone 3-dioxygenase, anthocyanidin synthase, chalcone isomerase, dihydroflavonol 4-reductase, shikimate O-hydroxycinnamoyltransferase, CYP98A/C3’H, caffeoyl-CoA O-methyltransferase, flavonoid 3-5-hydroxylase, flavone synthase I, UDP-glucose:3-O-d-glucosyltransferase, anthocyanidin 3-O-glucoside 2-o-xylosyltransferase and anthocyanidin 5,3-O-glucosyltransferase were identified with genome annotation (Table S11). These genes are involved in producing major anthocyanins such as cyanidin, delphinidin, pelargonidin, pelargonidin-3-glucoside, pelargonidin-3-sambubioside, cyanidin-3-glucoside, cyanidin-3-sambubioside, cyanidin-5-glucoside and cyanidin-3,5-diglucoside. Although ‘Kensington Pride’ and *M. laurina* mature fruits have yellow to orange and yellow colour skin, respectively, structural genes involved in anthocyanin biosynthesis were identified in the two genomes (Tables S12, 13). However, the total number of genes linked with anthocyanin biosynthesis was low (111 and 117 in ‘Kensington Pride’ and *M. laurina*, respectively) compared to that of ‘Irwin’ genome. Although in the *M. laurina* genome, structural genes for all 16 enzymes associated with anthocyanin biosynthesis were identified, genes only for 15 enzymes were identified in ‘Kensington Pride’ genome and gene/s encoding anthocyanidin 5,3-O-glucosyltransferase were not identified. Information on the number of genes encoding enzymes related to anthocyanin biosynthesis is summarized in Table S11, 12 and 13 for ‘Irwin’ and ‘Kensington Pride’ and *M. laurina* respectively. Anthocyanin biosynthesis is regulated by three major classes of transcription factors (TFs): MYB, bHLH, and WD40 proteins^[16]^. R2R3-MYB *MiMYB1* has been identified as the key MYB regulator in mango (cultivar ‘Irwin’) red coloration in fruit skin^[17]^. The *MiMYB1* gene sequence of the ‘Irwin’ and ‘Kensington Pride’ genomes was identical to that of the previously identified gene. The *M. laurina MiMYB1* gene had few SNPs, but they were not located in the R2, R3 domains or bHLH motif regions which are important conserved regions in the gene.

Carotenoids synthesized through the terpenoid pathways (carotenoid biosynthesis) are responsible for lighter (yellow to red) fruit skin and flesh color and β-carotene, lutein, and violaxanthin represent some of the major carotenoids identified in mango. In all three genomes, genes encoding all 13 enzymes which are associated with carotenoid biosynthesis were identified. Except for five genes (zeaxanthin epoxidase, phytoene synthase, lycopene epsilon-cyclase, phytoene desaturase, violaxanthin de-epoxidase) where the number of gene copies across the genomes varied by 1-2, all the other carotenoid biosynthesis genes had an identical number of gene copies in all three genomes (Table S14-16). A total of 28 genes were identified in ‘Kensington Pride’ and *M. laurina* while 23 genes were identified in ‘Irwin’.

### Volatile compounds synthesis genes

Mango fruits have high demand due to their taste and flavour, which are determined by volatile compounds produced in the fruit. Although the production of terpenoids, volatile compounds responsible for aroma and flavour, has been identified in mango fruits, the structural genes involved in terpenoid biosynthesis have not yet been characterized. The two *M. indica* genotypes, ‘Kensington Pride’ and ‘Irwin’, have high consumer preference and exhibit contrasting aroma profiles. Therefore, to analyze differences in the presence or absence and copy number of structural genes involved in terpenoid production, and to identify the genes responsible for the synthesis of unique terpenoids in two *M. indica* genotypes and *M. laurina*, the KEGG pathway analysis was conducted. Although the aroma profile of *M. laurina* has not yet been studied, we analyzed the structural genes associated with terpenoid biosynthesis to better understand the genetic basis of aroma-related traits in this wild relative. Here, key genes responsible for producing isopentenyl diphosphate (IPP) and dimethylallyl diphosphate (DMAPP); building blocks to produce terpenoids were identified in both *M. indica* cultivars and *M. laurina*. IPP and DMAPP are produced in two biosynthesis pathways, the mevalonate pathway and the non-mevalonate pathway or the MEP/DOXP pathway involving a series of enzymatic reactions (Figure 4). A total of 104, 106, and 108 genes are involved in IPP and DMAPP biosynthesis in ‘Irwin’, ‘Kensington Pride’ and *M. laurina*, respectively. Moreover, ‘Irwin’, ‘Kensington Pride’, and *M. laurina* were revealed to have 60, 72, and 80 genes linked to monoterpenoid biosynthesis (Figure 4, Table S17-19). Monoterpenoid synthase genes such as linalool Synthase, myrcene synthase, 4S-limonene synthase, α-terpineol synthase, and 1,8-cineole synthase were identified in all three genomes which involve producing (+)-linalool, myrcene, limonene, α-terpineol, and 1,8-cineole respectively. Among the three genomes, *M. laurina* had the highest number of gene copies for myrcene, limonene, α-terpineol, and 1,8-cineole biosynthesis (13,17,13, 17 genes respectively) whereas ‘Irwin’ had the highest number of genes for linalool biosynthesis (10) (Table S17-19). However, comparative analysis of the ‘Irwin’ and ‘Kensington Pride’ genomes revealed that Irwin had a higher number of genes associated with linalool, myrcene, and α-terpineol biosynthesis, whereas Kensington Pride exhibited a higher number of genes for limonene and 1,8-cineole biosynthesis. In addition, 88 genes were associated with the diterpenoid biosynthesis pathway in ‘Irwin’ whereas in ‘Kensington Pride’ and *M. laurina*, 78 and 85 genes were identified, respectively. Although the diterpenoids produced in ‘Irwin’, ‘Kensington Pride’, and *M. laurina* were the same, the number of genes copies encoding the corresponding enzymes varied slightly among them. (Table S20-22). Compared to other two genomes, structural genes for four sesquiterpene synthases ((Z)-γ-bisabolene synthase, valencene synthase, vetispiradiene synthase, and (+)δ-cadinene synthase) and seven triterpenoid synthases(seco-amaryin synthase, isomultiflorenol synthase, tirucalladienol synthase, baruol synthase, thalianol synthase, arabidiol synthase, marneral synthase) were only identified in the ‘Kensington Pride’ genome (Table S23-25). Out of four ‘Kensington Pride’ specific sesquiterpenoid synthase genes, two gene copies were identified for valencene synthase and each of the other three sesquiterpenoid synthases had only one gene copy. A single structural gene was responsible for encoding all seven unique triterpenoid synthases identified in ‘Kensington Pride’ which had only one gene copy in the genome (Table S24). Although we didn’t identify ‘Irwin’ specific triterpenoid/sesquiterpenoid synthase genes compared to other two genomes, *M. laurina* had two species-specific sesquiterpene synthase genes (lupan-3-β-20-diol synthase, camelliol C synthase) among 83 genes linked with sesquiterpene and triterpene biosynthesis (Figure 4, Table S25).

**Figure 4:**
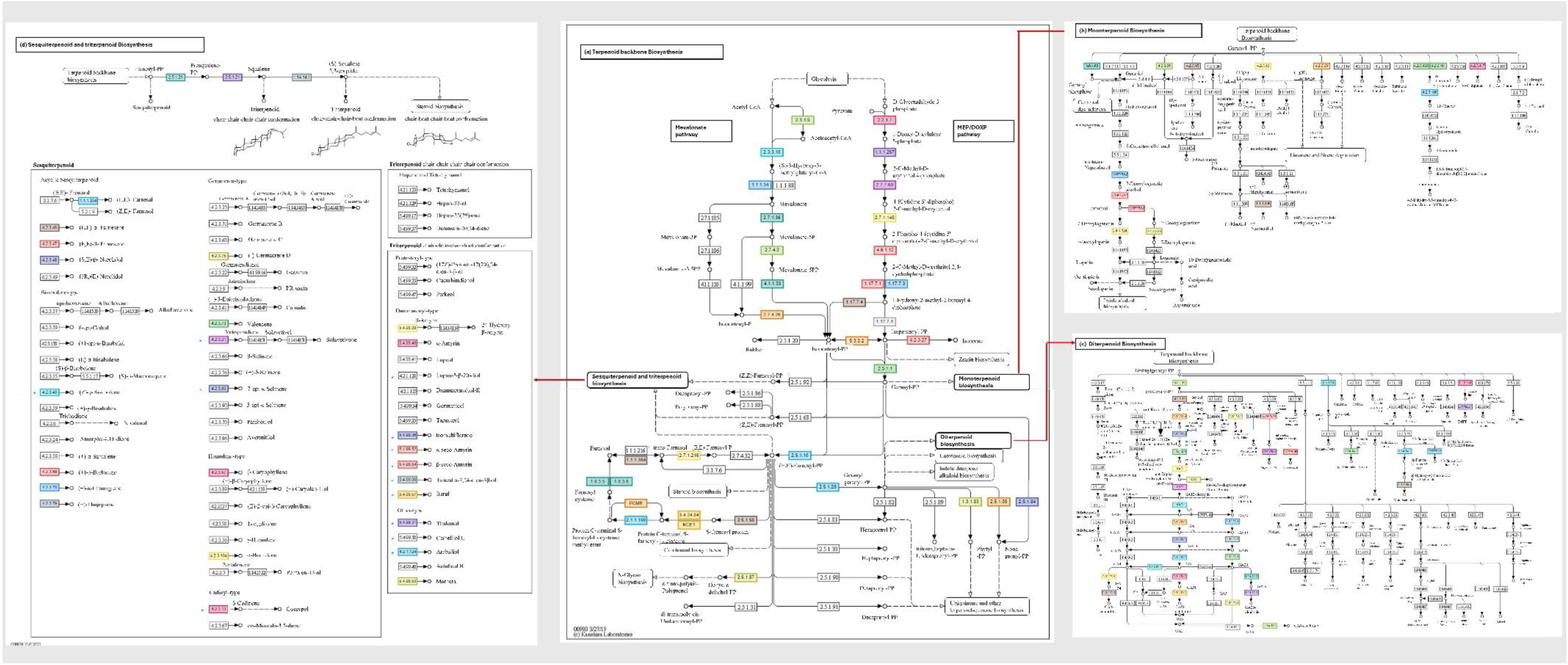
Terpenoid biosynthesis in ‘Kensington Pride’ mango. Here, biosynthesis pathways of ‘Kensington Pride’ are shown as representative for all three genomes. Coloured boxes indicate the enzymes which were identified by the annotation; therefore, the linked end-products are supposed be synthesized. For the boxes which are not coloured, associated enzymes are not identified by the genome annotation. All the different terpenoids are synthesized from two building blocks, isopentenyl diphosphate and dimethylallyl diphosphate which are produced by two different pathways in (a)Terpenoid backbone biosynthesis; mevalonate pathway and the non-mevalonate or MEP/DOXP pathway respectively. Then the enzymatic reactions of prenyltransferases synthesize higher-order building blocks such as geranyl diphosphate, and geranylgeranyl diphosphate and farsenyl diphosphate, which act as precursors for (b) monoterpenoid (C10), (c) diterpenoid (C20), and (d) sesquiterpenoid (C15) biosynthesis respectively. Although the number of genes associated with mono and diterpenoid biosynthesis were different among ‘Irwin’, ‘Kensington Pride’ and, *M. laurina*, the end products thought to be synthesized were same in all three genomes. However, comparison of sesquiterpene and triterpenoid biosynthesis pathways revealed unique tri and sesquiterpenes identified in ‘Kensington Pride’ and *M. laurina* which are indicated respectively in blue and red asterisk symbols in the figure.

### Key volatile compounds in ‘Kensington Pride’ and ‘Irwin’ mango pulp

In previous studies, volatile profiles of mango and their variation among different cultivars have been assessed with solvent extraction and solid phase microextraction methods. However, the production of volatile compounds in cultivars has not been confirmed with the presence of the respective structural genes in their genomes. Here we selected five key aroma volatile compounds (α-terpinolene, D-limonene, ꞵ-myrcene, 2-carene, 3-carene, and α-pinene) present in mangoes, which contribute significantly to aroma and flavour, and checked the presence of these compounds in the two cultivated mango genotypes while confirming the presence of the structural genes in the two respective genomes. All five volatile compounds were identified both in ‘Kensington Pride’ and ‘Irwin’ with the Head-space Solid-phase microextraction (HS-SPME)/ gas chromatography-mass spectrometry (GC-MS) method. It confirmed their production in these two genotypes by determining the retention time and peak areas of the spectra (Tables S26, and S27). The peak areas of the spectra is positively correlated with the concentration of the volatile compound identified in the genotypes. The highest peak areas in ‘Kensington Pride’ and ‘Irwin’ were obtained for the α-terpinolene and 3-carene, respectively, having significantly high peak area values compared to those of other volatile compounds. Out of the five volatile compounds identified using HS-SPME/ GC-MS in this study, genes encoding enzymes to produce D-limonene and ꞵ-myrcene were identified by KEGG pathway analysis in both genotypes.

## Discussion

The availability of high-quality genomes is vital for crop genetic studies and advancing molecular breeding. In this study, we assembled genomes for the most widely cultivated Australian mango cultivar, ‘Kensington Pride’, and the wild species *M. laurina*. To date, PacBio HiFi sequencing and Oxford Nanopore technology, complemented by long-range data (Hi-C, optical mapping, trio data) have been used to develop highly contiguous genomes for plant species. However, our recently published genome for the mango cultivar ‘Irwin’^[11]^ showed the feasibility of developing high-quality genomes solely with HiFi data.

The high abundance of repetitive sequences in eukaryotic genomes is the major factor complicating genome assemblies. While most of the interspersed and tandem repeats can be spanned by long reads, the assembly of satellite repeats; a type of extra-long tandem repeat, remains challenging due to the difficulty in spanning an entire satellite with long reads^[18]^. With PacBio HiFi reads at high genome coverage, we were able to assemble high-quality genomes for ‘Kensington Pride’ and *M. laurina* with 100% assembly completeness and high contig N50 values (15.1 Mb and 15.9 Mb, respectively). The quality of these genomes is comparable to the recently published genome for ‘Irwin’^[11]^, which has the highest completeness and contiguity among currently available genomes^[13,19,20]^. All the chromosomes in the collapsed genomes of ‘Kensington Pride’ and *M. laurina* were assembled telomere-to-telomere, containing few gaps (five and four gaps, respectively). Most of these gaps corresponded to ribosomal DNA clusters organized as long tandem arrays, which have been identified as one of the most challenging regions in the genomes to assemble^[21]^. Although the haplotype assemblies are less contiguous compared to collapsed genomes, they had 99.8-100% assembly completeness having almost all the telomeres. Here, the genomes assembled for ‘Kensington Pride’ and *M. laurina* further confirm that deep-sequenced, highly accurate HiFi reads alone can enable the assembly of near telomere-to-telomere genomes.

The genome size of angiosperms varies enormously. Among the different mechanisms influencing genome size, such as the tandem repeats, transposable elements, and polyploidization, repeated DNA sequences account for the majority of the genome size variations^[22]^. While tandem repeats generally contribute to a smaller proportion of the genome, the main repetitive sequences are TEs of which LTR elements occupy the largest proportion^[23]^. Among the three mango genomes studied, *M. laurina* had the largest genome size, followed by the two *M. indica* cultivars. Accordingly, *M. laurina* had the highest repetitive content, followed by ‘Kensington Pride’ and ‘Irwin’. In all three genomes, most of the repetitive sequences were unclassified repeats. This may be due to the presence of new TEs not classified by the tool or an under-representation of mango or closely related taxonomic groups in the repeat element reference database. Among the classified repeats, LTR elements, which have been identified as the most prevalent TEs in many other plant genomes, including peach, tomato, and maize^[24]^, occupied the largest proportion in all three genomes. Achieving 98.6-99.5% annotation BUSCO in all collapsed and haplotype assemblies of ‘Kensington Pride’ and *M. laurina* further confirmed the completeness of all the genomes.

In many plants, self-incompatibilities (outcrossing) and distant hybridization are the main reasons for high genome heterozygosity. Mangoes are generally heterozygous, and analysis of 22 cultivated mangoes in China has revealed high levels of heterozygosity^[13]^. Our results for ‘Kensington Pride’, and the recently published ‘Irwin’^[11]^ genome, also confirm that cultivated mangoes are highly heterozygous. Furthermore, reporting considerably higher heterozygosity for *M. laurina* (2.22%) compared to cultivated mango suggests that wild mango relatives are more heterozygous, a pattern previously observed in many other plant species, including cereals, legumes, and oil crops^[25]^. Genome synteny analysis identified local structural variations among the three genomes, with some structural variations unique to *M. laurina*. Structural variations, including insertions, deletions, duplications, and inversions, are identified as causative genetic variants for many traits related to crop domestication, improvement, and modern breeding. For example, the wild progenitor of cultivated tomatoes was discovered with numerous structural variations, many of which are associated with genes regulating fruit quality^[26]^. Furthermore, a link between chromosomal inversions and variations in breeding traits has been identified in cultivated mangoes^[27]^. Therefore, future population studies on *M. laurina* and other wild relatives, exploring polygenic traits to which structural variations are linked, will be useful in selecting progenies with desired traits in mango breeding programs. In addition, recent development of pangenomes with haplotype assemblies has enhanced the accuracy of identifying heterozygosity and structural variations in plants. For instance, a phased pangenome developed for potato with 60 haplotypes, including cultivated and wild relatives has identified the evidence of transposable elements in generating the structural variants and enhanced heterozygosity in cultivated potato compared to that of wild relatives ^[28]^. An extensive genetic variation also has identified in moso bamboo haplotype-based pangenome assembled with 16 accessions developed ^[29]^. This evidence suggests that, due to the high heterozygosity of mango, developing pangenomes with haplotype-resolved assemblies would allow for high-resolution identification of structural variations in the future.

Throughout the evolutionary history of angiosperms, multiple polyploidization/WGD events have been uncovered, leading toward complexities and novelties in the genomes enhancing the species diversification, reproductive isolation, and environmental adaptation. All core eudicots share one genome triplication event in their evolutionary history, while many species-rich angiosperm families display evidence for more rounds of ancient polyploidization^[30]^. Wang et al^[13]^ identified that a recent WGD event occurred in mango (family *Anacardiaceae*), approximately 33 MYA, after it diverged from the *Rutaceae* and *Sapindaceae* families (∼ 70 MYA). Our study also confirms that cultivated mango shows these two WGD events.

Analysis of the origin of duplicated genes in a species could further provide evidence for polyploidization events. The number of segmental/WGD duplicates in a genome depends on the number of polyploidization events that occurred, their timing, and the level of gene retention after each WGD event^[31]^. According to a recent study, *Vitis vinifera* (grapes) has undergone only one polyploidization, which is common to all the eudicots (more than 100 MYA)^[32]^ and has 15% segmental/WGD genes. In contrast, *Populus trichocarpa* and *Glycine max*, which have undergone one and two additional lineage-specific WGD events, respectively, have 51.6% and 76.0% segmental/WGD duplicated genes^[31]^. Therefore, classifying more than 50% of the genes under segmental/WGD duplicates further supports that mango has undergone an additional lineage-specific WGD event. Many plant species that have only undergone the WGD event common to all eudicots have dispersed duplicates as the highest number of duplicated genes. In contrast, mango and many other species that have at least one additional WGD, have dispersed duplicates as the second most abundant type of duplicated genes, with tandem and proximal duplicates in smaller proportions^[31]^. Collinearity analysis revealed a high level of synteny and collinearity among all three mango genomes. The presence of a significant level of rearrangements in collinear blocks, including duplications and translocations, further supports recent polyploidization. Considering the number of collinear genes shared among the three genomes, ‘Irwin’ and ‘Kensington Pride’ shared a slightly smaller number of collinear genes compared to genes shared between ‘Irwin’ and *M. laurina*. According to evolutionary relationships of the genus, *M. laurina* has been identified as a species that exhibits a distinct chloroplast genome, with a close evolutionary relationship with domesticated mango ^[33]^. However, no strong or conclusive evidence has yet been found to support the occurrence of hybridization between the two species. Therefore, less co-linearity between ‘Irwin’ and ‘Kensington Pride’ could be due to local rearrangements in the ‘Kensington Pride’ and ‘Irwin’ genomes, such as inversions, insertions, or deletions, which could either alter the gene order and positions, thereby reducing the number of collinear genes between the two genomes.

Anthracnose is one of the most serious diseases affecting mango, causing a 30-60% loss in fruit yield, which can reach up to 100% under humid environmental conditions^[34]^. A recent transcriptomic study analyzed plant responses after infection of mango fruits with *C. gloeosporioides*^[35]^. The results identified 35 upregulated defense-related genes, including ethylene response factors, nucleotide binding site-leucine-rich repeats, nonexpressor of pathogenesis-related genes and pathogenesis-related proteins (6). However, the results did not identify specific genes involved in anthracnose resistance in mangoes. Felipe et al^[15]^ identified that a SNP in the *β-1,3-GLU2* gene enhances the anthracnose resistance by hydrolyzing the fungal cell wall. However, our results indicate the presence of multiple copies of *β-1,3-GLU2* genes, with both the resistant and susceptible SNPs present in the susceptible ‘Irwin’, and moderately suseptible ‘Kensington Pride’ as well as the resistant *M. laurina*. Transcriptomic studies in other species, including lupin^[36]^ and an anthracnose-resistant genotype of bean^[37]^, have identified over-expression of β-1,3-glucanases after inoculation with *Colletotrichum* sp. suggesting their importance in anthracnose resistance. Even though the *β-1,3-GLU2* gene has been suggested to enhance anthracnose resistance in mango, our results indicate the need for gene expression analysis to validate its role in disease resistance by confirming the differential expression of the genes in *M. laurina*, ‘Irwin’ and ‘Kensington Pride’.

Fruit skin color is an important trait in mango that significantly influences consumer preference. Anthocyanins, responsible for red coloration, are produced via the flavonoid biosynthesis pathway. A recent study analysed anthocyanin levels produced in mango cultivars with different peel colors at the ripening stage. According to their results, a greater concentration of anthocyanins, specifically cyanidin-3-O-glucosides and peonidin-3-O-glucosides, was found in red peel cultivar compared to green and yellow peel cultivars^[5]^. Furthermore, higher gene expression levels have been observed for the selected genes related to anthocyanin biosynthesis in red peel cultivar, whereas cultivars with green and yellow colored peel have shown relatively lower expression levels. In this study, we identified functionally characterized genes involved in anthocyanin biosynthesis in all three mango genomes. The higher number of structural genes identified in the ‘Irwin’ genome compared to the other two genomes supports the view that more genes may have be involved in producing red pigmentation in ‘Irwin’ fruit skin. Among different transcription factors (MYB, bHLH, and WD40 proteins) regulating anthocyanin biosynthesis, the MYB transcription factor R2R3-MYB *MiMYB1* has shown a higher expression level in ‘Irwin’^[17]^. However, our results revealed that the gene sequences of conserved regulatory domains in *MiMYB1*were similar to that of ‘Kensington Pride’ and *M. laurina*. Therefore, future comparative gene expression analysis of *MiMYB1* and other TFs may provide a deeper understanding of the regulation of anthocyanin biosynthesis in mangoes with different peel colors.

Carotenoids are another group of pigments that give fruit peel their yellow-to-orange colour. Our results characterized the structural genes involved in carotenoid biosynthesis in all three genomes, including those for β-carotene, lutein, zeaxanthin, and violaxanthin. Karanjalker et al^[5]^ discovered that total carotenoid content in yellow-colored cultivars was higher compared to green and red-colored cultivars, revealing β-carotene and violaxanthin as the major compounds produced in peel. Furthermore, gene expression analysis has suggested that *lycopeneβ-cyclase* and *violaxanthinde-epoxidase* gene expression was positively correlated with β-carotene and violaxanthin content in fruit peel. Our results revealed that more genes are involved in carotenoid biosynthesis in ‘Kensington Pride’ and *M. laurina*, which exhibit yellow color peel at the ripening stage, compared to ‘Irwin’. Since all the structural genes related to carotenoid biosynthesis were characterized for ‘Irwin’, ‘Kensington Pride’, and *M. laurina*, these resources could be used in future studies to analyze gene expression, regulation, and inheritance patterns.

Mango’s high consumer preferences is mainly due to its distinctive flavour, resulting from a complex blend of aroma volatile compounds. Although hundreds of volatile compounds have been characterized including terpenes, esters, alcohols, aldehydes, ketones, fatty acids and lactones, terpene hydrocarbons (monoterpenes) have been identified as the most abundant group of volatile compounds in mango^[38–40]^. To date, genes involved in terpenoid biosynthesis have not been characterized in mangoes. Furthermore, no molecular markers have been developed specifically for fruit aroma volatile compound biosynthesis genes, which are useful in selecting progenies with desired traits in breeding. Here, we identified functionally annotated genes that encode terpenoids and validated some of these compounds by HS-SPME/ GC-MS method. The volatile profile of mango varies considerably with the cultivar^[41]^ and α-terpinolene is the key and most abundant volatile compound responsible for the characteristic flavour in ‘Kensington Pride’^[42]^. Although we identified the production of α-terpinolene in ‘Kensington Pride’, genes specifically encoding α-terpinolene were not identified since this compound is not included in the monoterpenoid biosynthesis pathway of KEGG analysis. However, among two different classes of terpene synthases present in plants, class I terpene synthases are capable of producing multiple terpenes from a single substrate^[43]^. In *Arabidopsis thaliana,* a monoterpene synthase has been revealed to produce 1,8-cineole as the main product along with nine minor monoterpenes, including terpinolene, α-terpineol, α-pinene, myrcene, sabinene, β-pinene, limonene, β-ocimene, and (+)-α-thujene^[44]^. Therefore, terpene synthase 10, and probable terpene synthase 12 genes, linked to monoterpenoid biosynthesis, could be potential candidate genes encoding α-terpinolene as well as 3-carene, 2-carene, and α-pinene in ‘Kensington Pride’, where the production of all these monoterpenes have been identified previously^[42]^ and confirmed in our study. Furthermore, among unique structural genes identified in ‘Kensington Pride’ encoding tri and sesquiterpenes, the presence of bisabolene in the fruit has been previously identified^[42]^. Future studies on the expression of these unique genes and identifying the encoded volatiles in the fruit (such as vetispiradiene, (+)-delta cadinene, seco-amaryin, isomultiflorenol, tirucalladienol, baruol, thalianol, arabidiol, and marneral), could provide a deeper understanding of their contribution to the unique flavour of ‘Kensington Pride’. Similar to ‘Kensington Pride’, specific genes encoding 3-carene, the main volatile compound identified in ‘Irwin’, were not identified. However, multiple copies of terpene synthase 10, and probable terpene synthase 12 genes identified in the genome might be responsible for 3-carene biosynthesis. Furthermore, though the volatile compounds in wild relatives have not been identified to date, we characterized the structural genes of the main volatiles produced in *M. laurina*, which included two unique genes encoding sesquiterpenes; lupan-3beta,20-diol, and camelliol C. Future research on the volatile profile of *M. laurina* will further facilitate their use in mango breeding.

The high-quality genomes we assembled for ‘Kensington Pride’ and the wild relative, *M. laurina,* comparative genome analysis together with the recently published ‘Irwin’ genome provide valuable insights into genes associated with fruit quality traits. Furthermore, the *M. laurina* genome is a valuable resource for analyzing gene expression associated with anthracnose resistance. Additionally, these genomes will facilitate the development of molecular markers for desired traits, thereby supporting advancements in mango breeding.

## Material and methods

### Plant materials, DNA extraction, and sequencing

Fresh young leaves of *M. indica* cv. ‘Kensington Pride’ and *M. laurina* were collected from trees located at the Walkamin Research Station, Mareeba, (17°08̍ 02″S and 145°25 37″E), North Queensland, Australia. Genomic DNA was extracted using a cetyltrimethylammonium bromide (CTAB) method^[45]^ with modified steps^[11]^. Extracted DNA was evaluated for quality and quantity. PacBio HiFi sequencing of the two species was performed each in two PacBio Sequel II SMRT cells at the Institute for Molecular Bioscience, The University of Queensland, Australia.

### RNA extraction and Illumina sequencing

Young leaf, flower buds, pre- and post-anthesis flower tissues of ‘Kensington Pride’ and *M. laurina* were collected from the trees at the Walkamin Research Station, Mareeba, North Queensland, Australia. RNA was extracted using a CTAB method^[46]^ with modifications and Qiagen RNeasy Mini Kit (Qiagen, Valencia, CA, United States) was used to purify the extracted RNA. Illumina short read sequencing was performed at the Australian Genome Research Facility, University of Queensland.

### Draft genome assembly

PacBio HiFi read quality was evaluated with SMRT Link v11.0. HiFi reads were assembled by the HiFiasm Denovo assembler^[47]^ with default settings to generate a collapsed assembly and two haplotypes. The quality and the contiguity of the assemblies were assessed using Benchmarking Universal Single-Copy Orthologs (BUSCO) with viridiplantae database (BUSCO v 5.4.6)^[48]^ and the Quality Assessment Tool v5.2.0^[49]^ respectively. K-mer analysis was performed in Jellyfish (v2.2.10)^[50]^ using Illumina short reads trimmed (0.01 quality limits) in CLC Genomic WorkBench (CLC-GWB). The results were further analyzed in GenomeScope v2.0^[51]^ to determine the genome heterozygosity.

### Assembly of pseudomolecules

Contig level assemblies of the two collapsed genomes were first aligned with the published *M indica* cv. ‘Irwin’ genome^[11]^ in GENIES^[52]^. The contigs were then sorted and re-oriented concerning the reference genome. Based on their alignment with the reference, contigs were assigned to chromosomes. Telomeres in contigs were identified using TIDK v0.2.1 (https://github.com/tolkit/telomeric-identifier). The presence of telomeres at both ends of the contigs confirmed its representation of a single pseudomolecule. When more than one contig was assigned to a chromosome in which only one end had telomeric repeats or both ends didn’t have telomeric repeats, the nucleotide sequence was confirmed with NCBI nucleotide Blast. Then those contigs were linked by adding 100 N’s in between to imply that the two contigs were joined.

The two contig-level haplotype assemblies of ‘Kensington Pride’ and *M. laurina* were aligned with their respective collapsed genomes and contigs aligned with 20 chromosomes were identified. Once contigs were characterized based on the presence of telomeres or repetitive sequences at the ends, relevant contigs were joined to obtain complete chromosomes.

### Genome annotation, collinearity and WGD analysis

Collapsed genome and two haplotypes of ‘Kensington Pride’ and *M. laurina* were annotated structurally and functionally. Repetitive sequences were identified with Repeatmodeler2 v2.0.4^[53]^ and masked with Repeatmasker v.4.1.5^[54]^. HISAT2 tool^[55]^ was used to align quality and adapter trimmed RNA reads to the masked genome and structural annotation was performed using Braker3 v.3.0.3^[56,57]^. Omicsbox 3.0.30 ^[58]^ was used for functional annotation. A coding potential analysis was conducted for the CDS sequences that didn’t have blast hits during genome annotation using the already built model *Arabidopsis thaliana* and the model created for *M. indica*. The structural genes linked with important biosynthesis pathways, including carotenoid, anthocyanin, and terpenoid biosynthesis, were identified with KEGG pathway analysis^[59]^ in omicsbox v.3.0.30.

Inter-genome collinear blocks were determined by MCScanX^[29]^. Gene duplication analysis was determined using the duplicate_gene_classifier implemented in the MCScanX package. WGD events of the genomes were analyzed using ks distribution in WGDI^[60]^.

### Structural variant identification and orthologous cluster analysis

Collapsed genomes of ‘Kensington Pride’, ‘Irwin’^[11]^, *M. laurina*, and their haplotype assemblies were aligned pairwise using the MUMer software^[61]^. With the use of the delta filter implemented in Mummer, the alignments were filtered and the structural variations were analyzed using the SyRI tool^[12]^. Finally, the results were visualized using plotsr^[62]^. The unique gene clusters in ‘Kensington Pride’, ‘Irwin’ and *M. laurina* genomes were identified by clustering protein sequences of the genomes at e-value of 1e-2 with OrthoFinder algorithm in OrthoVenn3^[63]^. After extracting unique genes in unique gene clusters of the genomes, KEGG pathway analysis^[59]^ was conducted to identify genes related to important biological processes, cellular processes and key biosynthesis pathways.

### Anthracnose resistance gene analysis

In *β-1,3-GLU2* gene, region of gene sequence which include the SNP related to anthracnose resistance was identified^[15]^. Annotated genes for *β-1,3-GLU2* were extracted from ‘Kensington Pride’, ‘Irwin’ and *M. laurina* and aligned in Clone Manager Professional 9 to analyze the presence of resistant (Adenine) or susceptible (Guanine) SNP in the genes. Structural differences of the genes were identified using CLC-GWB.

### Aroma volatile compound analysis in ‘Kensington Pride’ and ‘Irwin’

#### Mango fruits

’Irwin’ and ‘Kensington Pride’ mango fruits were collected at commercial maturity from Southedge Research Station, Mareeba, Australia (16°45′S, 145°16′E). Fruits were stored at 10°C until they were ripe. Three biological fruit replicates each with two technical replicates were used for the two cultivars. For each biological replicate, three fruits were subsampled and one cheek from the flesh of each fruit was cut off. The cubed flesh for each replicate was pureed, dispensed into two glass vials and frozen at −80°C. Prior to instrumental analysis, samples were thawed from −80°C to −19°C overnight and then at room temperature. Pureed flesh was then blended with a stainless-steel blender and transferred back into glass vials.

### Head-space sampling and instrumental analysis

All the solvents used were HPLC grade, and all reagents and standards were purchased from Sigma-Aldrich, Australia. For each technical replicate of ‘Irwin’ and ‘Kensington Pride’, 3.5 g homogenized mango flesh was added to a 20 ml SPME vial (Merk, Australia) containing 3.5 ml of saturated sodium chloride solution and a magnetic stirrer flea (15x 4.5 mm). The vials were sealed with a rubber septum and 10 µl of combined internal standard solution was injected through the septum using a glass syringe at concentrations of 0.05mg/L for each of hexanoate, tridecane, and hexadecane. The content of the vial was heated to 40°C with stirring at 250 rpm for 2 mins. Extraction was performed with a grey (divinylbenzene/carboxen/polymethylsiloxane, 1 cm) fibre (Supelo/ USA), exposing to the headspace for 30 mins. The fibre was desorbed at 200°C for 8 mins by injecting it into a temperature programmable vaporizing inlet.

Samples were analyzed with an Agilent gas chromatograph (Agilent Technologies, USA) equipped with a Gerstel MPS2XL multi-purpose sampler and 5975N mass selective detector. The data were analyzed by MSD Chemstation E 02.021431 software. Separation was achieved in a DB-WAX capillary column (30 m x 0.25 mm) with 0.25 µm film thickness. Helium was used as the carrier gas with an average velocity of 44 cm/sec, a constant flow rate of 1.5 mL/ min while the pressure and the total flow were 75.7 kPa and 70.6 mL/ min respectively. The oven temperature was maintained at 40°C for 3 mins, followed by an increase to 120°C at 8°C/min and then to 220°C at 10°C/min which was held for 12 mins. The temperature of the mass spectrometer quadrupole temperature was set at 150°C and the source was set to 250°C. Ion electron impact spectra for selected volatile compounds were recorded with scan (35-350 m/z) mode. The target volatiles were identified by comparing their retention time with authentic compounds. Compound presence or absence was determined by presence or absence of a peak by this method. Internal standards were used to ensure reproducible SPME results run to run.

## Accession numbers

All the raw sequencing reads are deposited in the National Centre for Biotechnology Information (NCBI) under BioProject: PRJNA1148201. Mango genomes are deposited in the Genome Warehouse under Bioprojects PRJCA029779 and PRJCA029972.

## Acknowledgments

The authors would like to acknowledge the University of Queensland Research Computing Centre for the computational resources provided and Dr. Ian Bally for providing images of the ‘Kensington Pride’, and *M. laurina*.

## Supporting Information

**Figure S1**: K-mer profile (K=17) spectrums generated for three mango samples from Illumina sequence data using Genome scope. **Figure S2:** Genome alignments of Irwin, Kensington Pride, and *M. laurina***. Figure S3:** Summary of functional annotation for collapsed, hap1, and hap2 genomes of (a) Kensington Pride and (b) *M. laurina.* **Figure S4:** The coding potential assessment of CDS sequences that did not give a blast hit during functional annotations**. Figure S5:** Chromosome-wise structural variations (a) among Kensington Pride, Irwin and *M. laurina* genomes, between haplotypes of (b)*M. indica* Kensington Pride and (c) *M. laurina* genomes. **Figure S6:** Ks distribution peaks for paralogous gene pairs of Irwin, Kensington Pride, *M. laurina* and *Citrus sinensis.* **Figure S7:** Functions of unique genes in (a) Irwin, (b) Kensington Pride, and (c) *M. laurina* with related to biological process and molecular function and cellular component. **Figure S8:** Alignment of the region in β-1,3-glucanase 2 genes where the SNP for the anthracnose resistance is located. **Figure S9:** Anthocyanin biosynthetic pathway of Irwin as a representative of all three genomes. **Table S1:** Summary of sequence data generated by two PaBio SMRT cells. **Table S2:** Details of the repetitive sequences present at the ends of contigs which required joining to obtain complete pesudomolecules. **Table S3**: Details of contigs in the Kensington Pride collapsed genome and haplotypes**. Table S4**: Details of contigs in the *M. laurrina* collapsed genome and haplotypes. **Table S5**: Repetitive elements in the collapsed genome and two haplotypes of Kensington Pride. **Table S6**: Repetitive elements in the collapsed genome and two haplotypes of *M. laurina.* **Table S7**: Structural variations among Kensington Pride, *M. laurina* and published Irwin genomes. **Table S8**: Structural variations between haplotypes of Kensington Pride and *M. laurina* genomes. **Table S9**: Biosynthesis pathways associated with unique genes in three mango genomes. **Table S10**: Details of β-1,3-glucanase 2 gene copies in Kensington Pride, Irwin and *M. laurina* genomes. **Table S11**: Anthocyanin biosynthesis genes in Irwin. **Table S12** Anthocyanin biosynthesis genes in Kensington Pride. **Table S13**: Anthocyanin biosynthesis genes in *M. laurina*. **Table S14:** Genes related to carotenoid biosynthesis in Irwin. **Table S15**: Genes related to carotenoid biosynthesis in Kensington Pride. **Table S16**: Genes related to carotenoid biosynthesis in *M. laurina*. **Table S17** Genes related to monoterpenoid biosynthesis in Irwin. **Table S18** Genes related to monoterpenoid biosynthesis in Kensington Pride. **Table S19** Genes related to monoterpenoid biosynthesis in *M. laurina.* **Table S20** Genes related to diterpenoid biosynthesis in Irwin. **Table S21** Genes related to diterpenoid biosynthesis in Kensington Pride. **Table S22** Genes related to diterpenoid biosynthesis in *M. laurina*. **Table S23** Genes related to triterpenoid and sesquiterpenoid biosynthesis in Irwin. **Table S24** Genes related to triterpenoid and sesquiterpenoid biosynthesis in Kensington Pride. **Table S25** Genes related to triterpenoid and sesquiterpenoid biosynthesis in *M. laurina*. **Table S26** Identified terpenoids from ripe ‘Kensington Pride’ and ‘Irwin’ fruits. **Table S27**: Average peak areas for identified terpenoids from ripe Kensington Pride and Irwin fruits.

## Authors’ contributions

Study conception and design: RJH, AF, NLD; funding acquisition and project administration: RJH; Sample and data collection: UKW, AF, NLD, HES; formal analysis: UKW, RJH, AF, AKM, HES; Original draft preparation: UKW; review and editing of the original draft: RJH, AF, NLD, AKM, HES, UKW. All authors reviewed and approved the manuscript.

