## Supplementary material for "Chromosome-scale haplotype genome assemblies for the Australian mango ‘Kensington Pride’ and a wild relative, *Mangifera laurina*, provide insights into anthracnose-resistance and volatile compound biosynthesis genes": Figures: Supporting Information_Figures.docx

**Supplemental Figures**

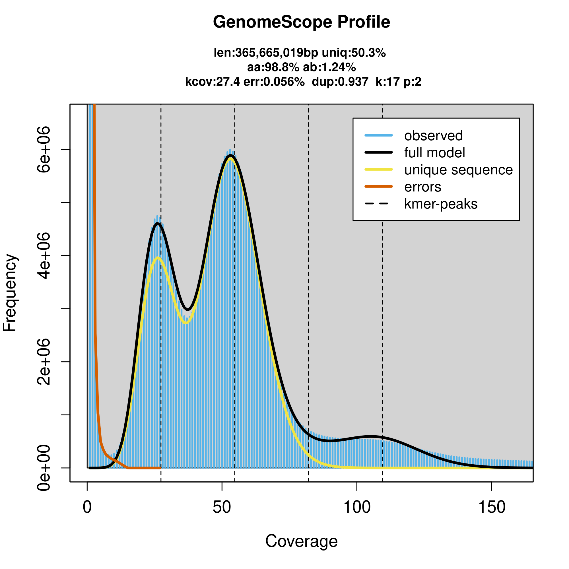

(a)

(b)
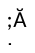
)

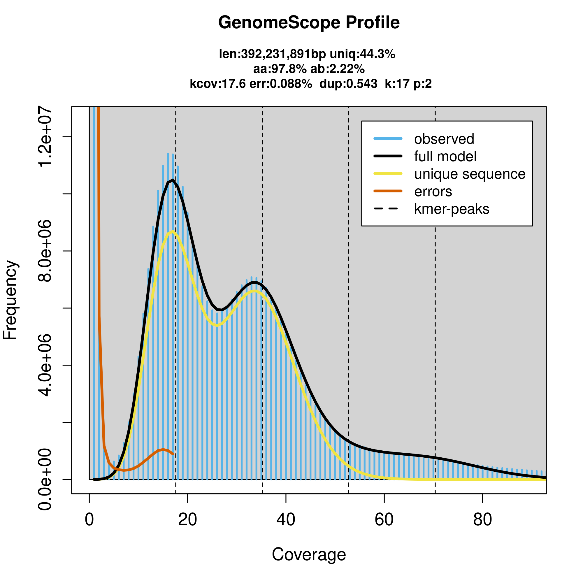

(c)

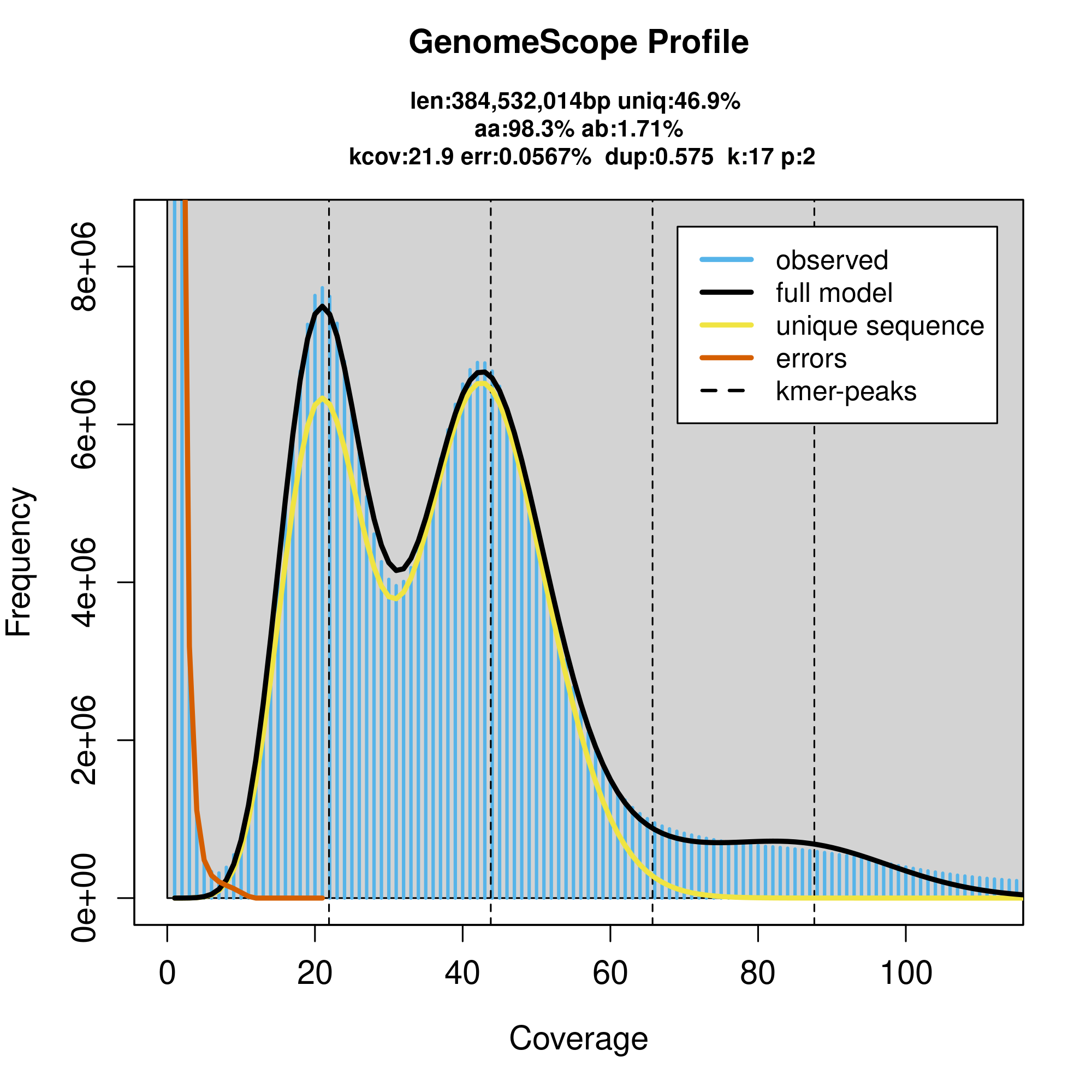

**Figure S1: K-mer profile (K=17) spectrums generated for three mango samples from Illumina sequence data using Genome scope.** (a) *M. indica* cv. Irwin, (b) *M. indica* cv. Kensington Pride, (c): *M. Laurina.* Bimodal profiles are characteristic of heterozygous genomes, while simple poisson profiles are characteristic of homozygous genomes. Here, the characteristic bimodal profile suggests that all three genomes are heterozygous.

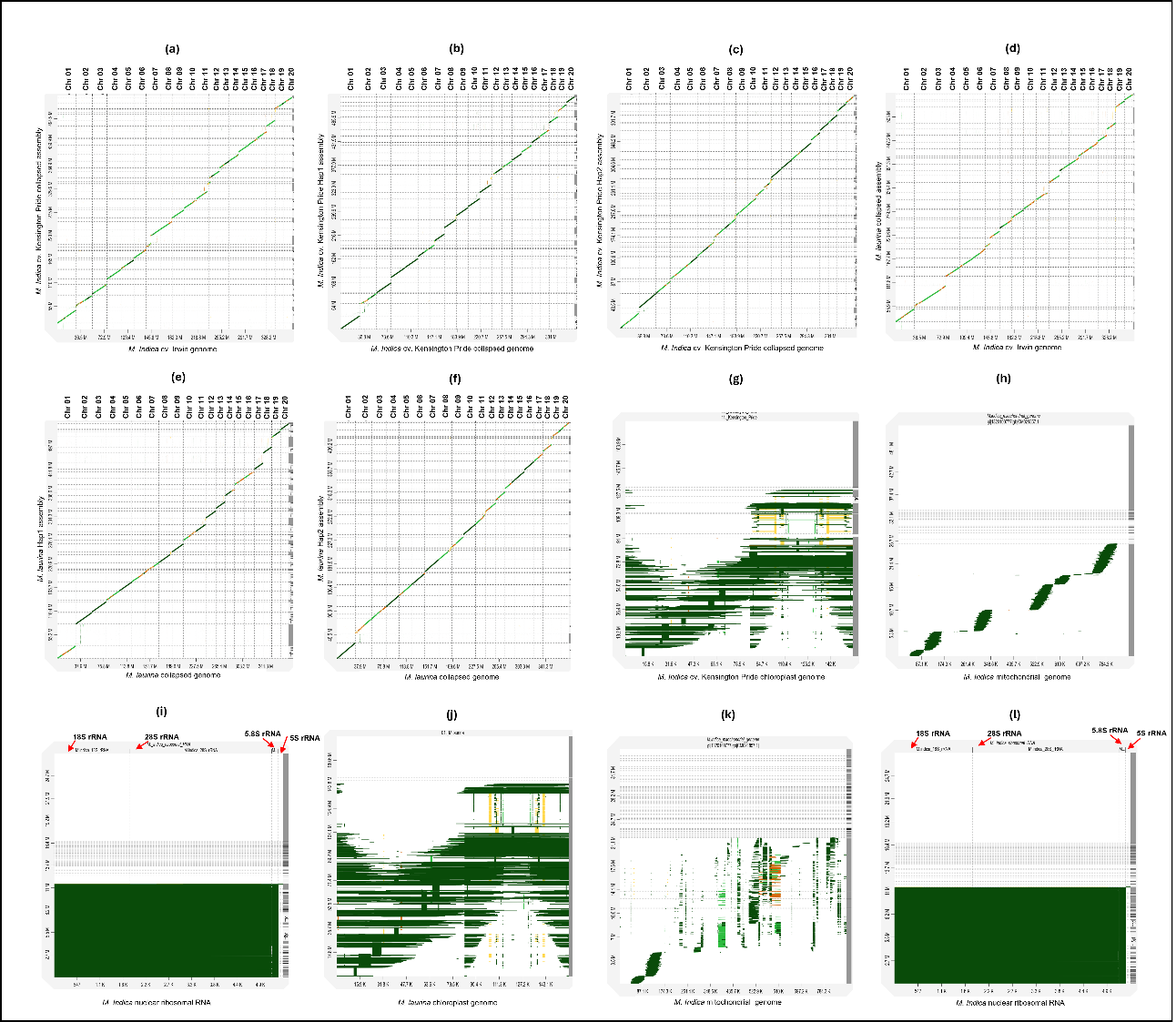
**Figure S2 Genome alignments of Irwin, Kensington Pride, and *M. laurina*.** (a) 'Irwin' genome vs 'Kensington Pride' collapsed assembly, (b) 'Kensington Pride' collapsed genome (20 chromosomes) vs 'Kensington Pride' hap1 assembly, (c) 'Kensington Pride' collapsed genome (20 chromosomes) vs 'Kensington Pride' hap2 assembly, (d) 'Irwin' genome vs *M. laurina* collapsed assembly, (e) *M. laurina* genome (20 chromosomes) vs *M. laurina* hap1 assembly, (f) *M. laurina* genome (20 chromosomes) vs *M. laurina* hap2 assembly.

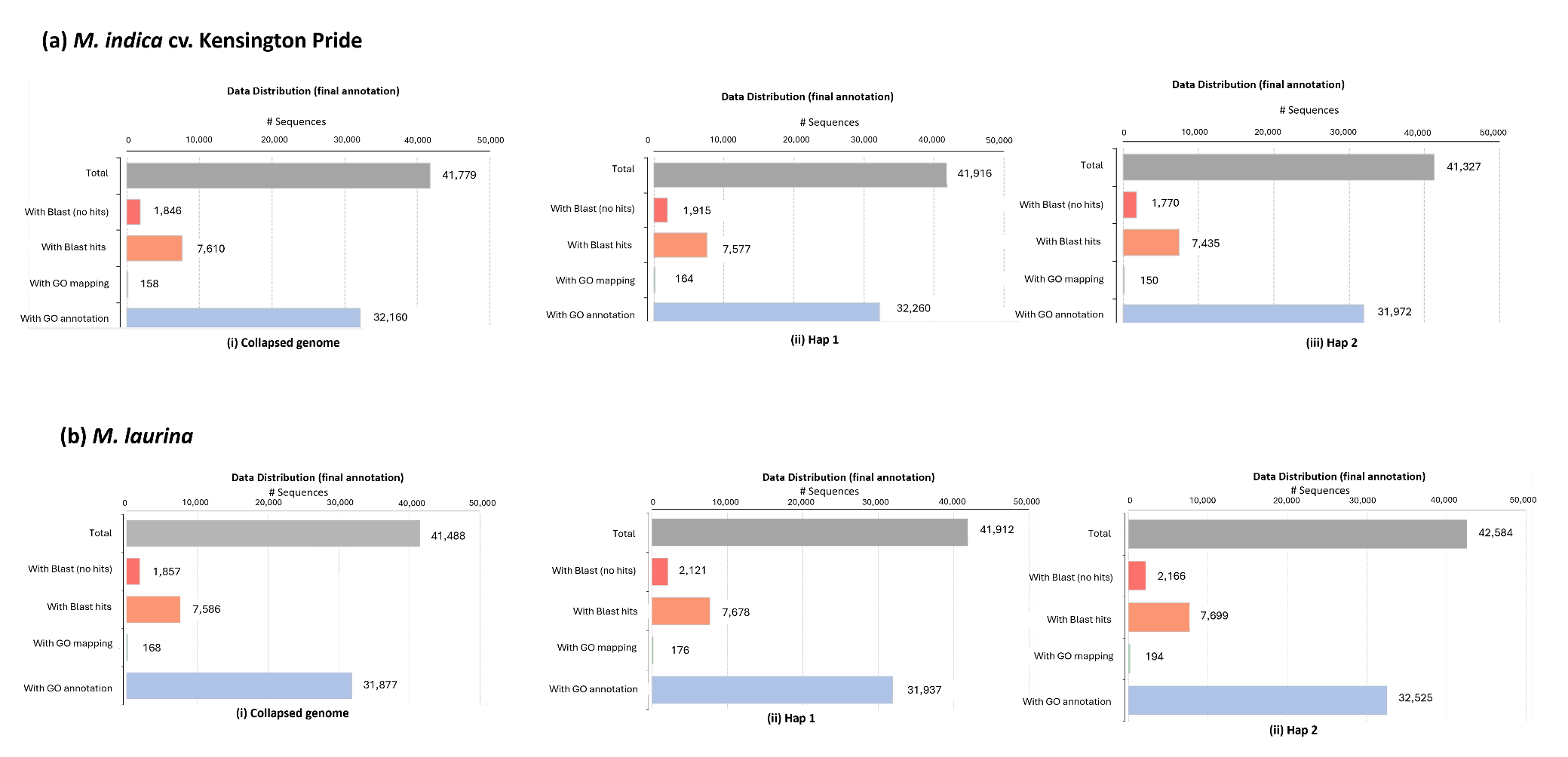

**Figure S3 Summary of functional annotation for collapsed, hap1, and hap2 genomes of (a) Kensington Pride and (b) *M. laurina***

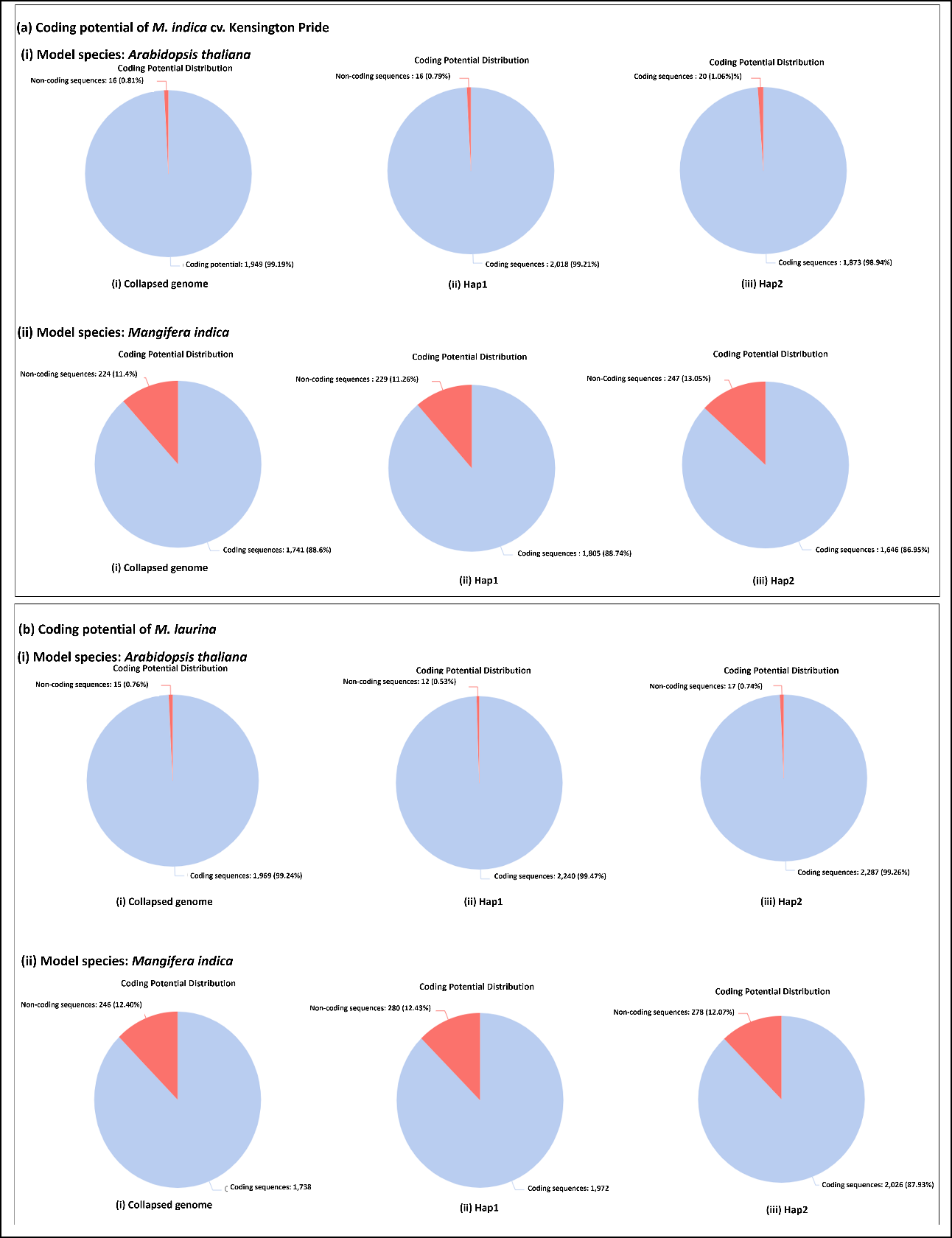

**Figure S4 The coding potential assessment of CDS sequences that did not give a blast hit during functional annotations. (**a) Kensington Pride (b)  *M. laurina.*  The analysis was conducted using (i) *Arabidopsis thaliana,* and (ii) *Manifera indica* as model species for the collapsed genome, hap1, and hap2.

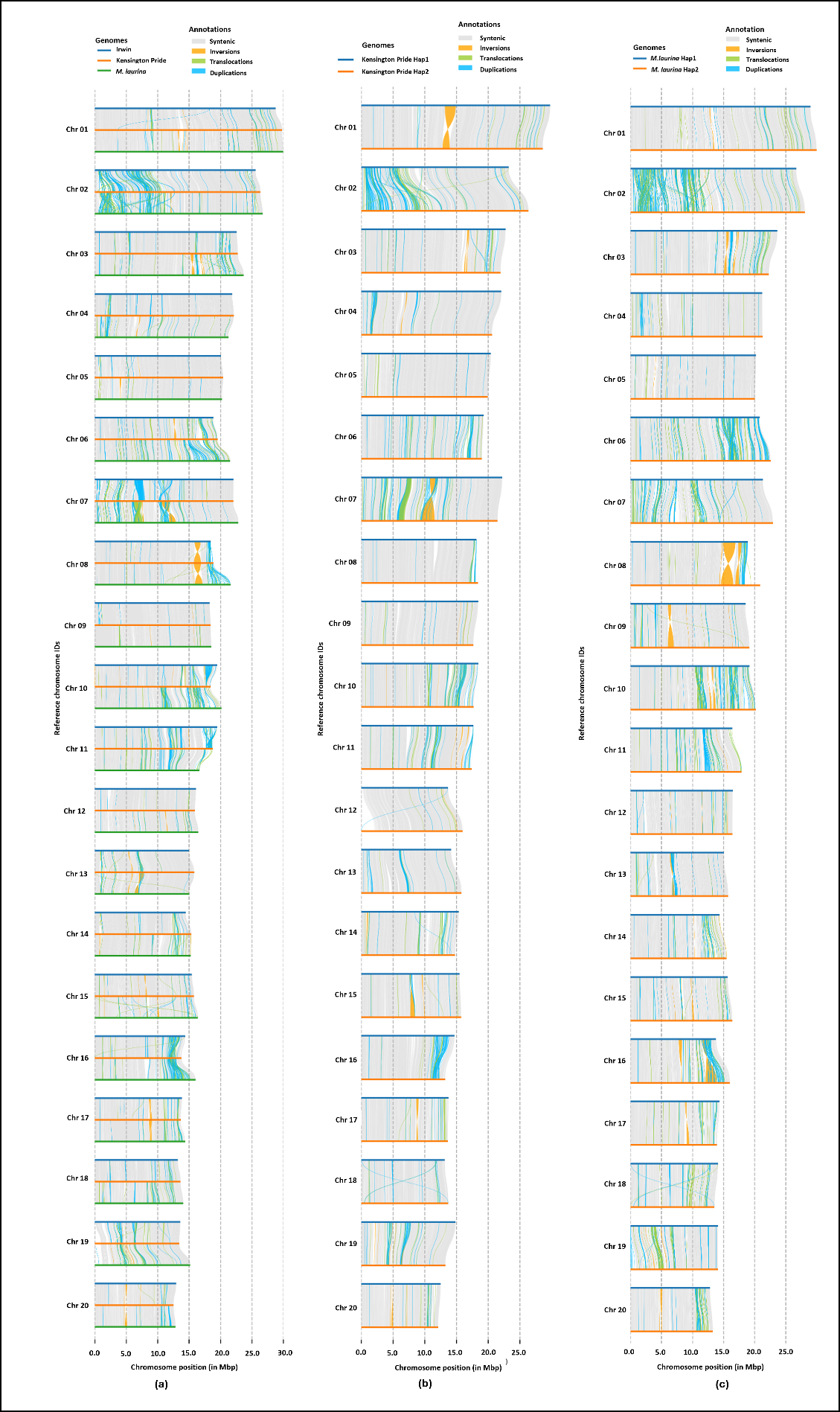
**Figure S5 Chromosome-wise structural variations (a) among Kensington Pride, Irwin and *M. laurina* genomes, between haplotypes of (b)*M. indica* Kensington Pride and (c) *M. laurina* genomes.**

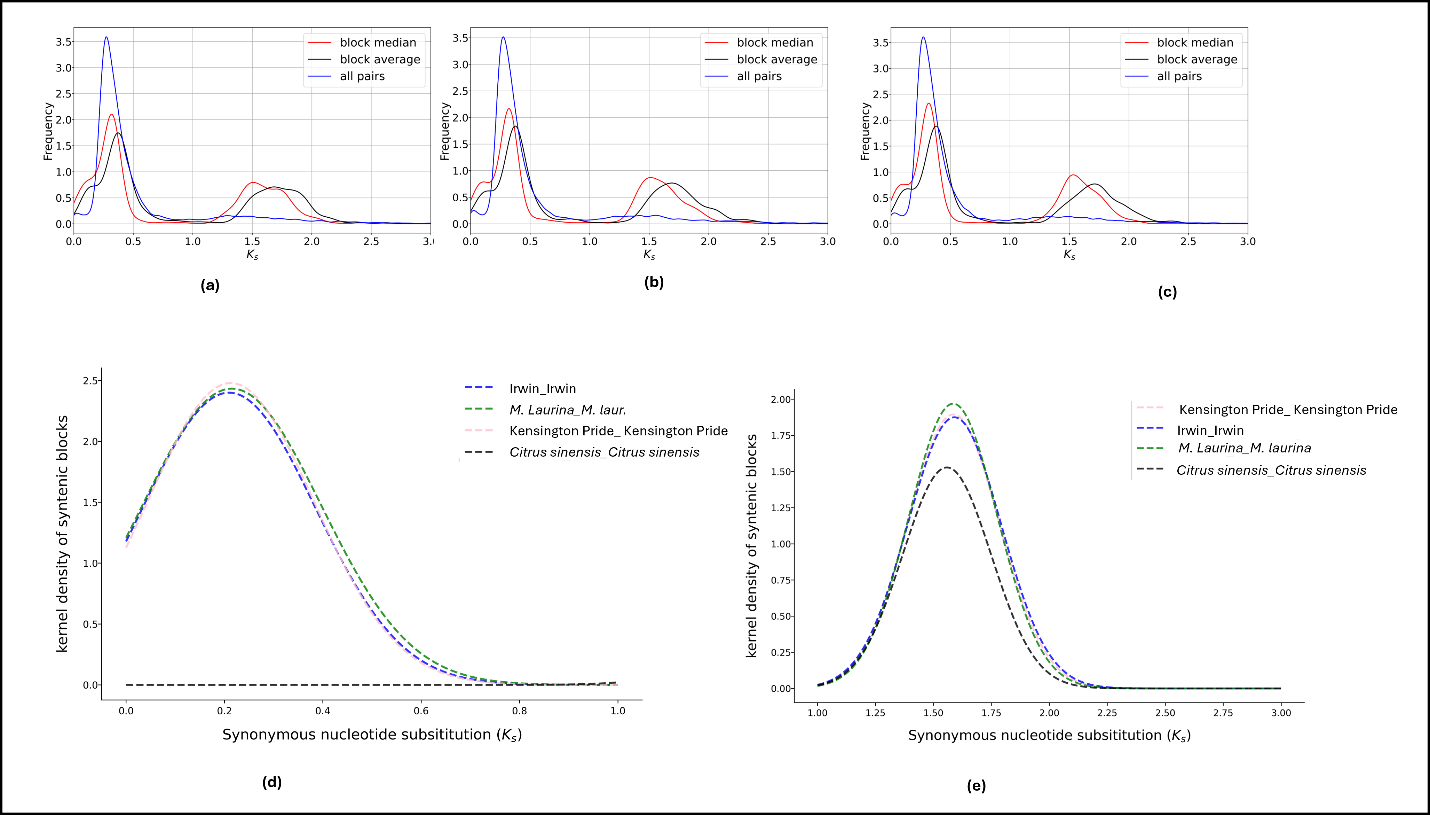
**Figure S6 Ks distribution peaks for paralogous gene pairs of Irwin, Kensington Pride, *M. laurina* and *Citrus sinensis.***

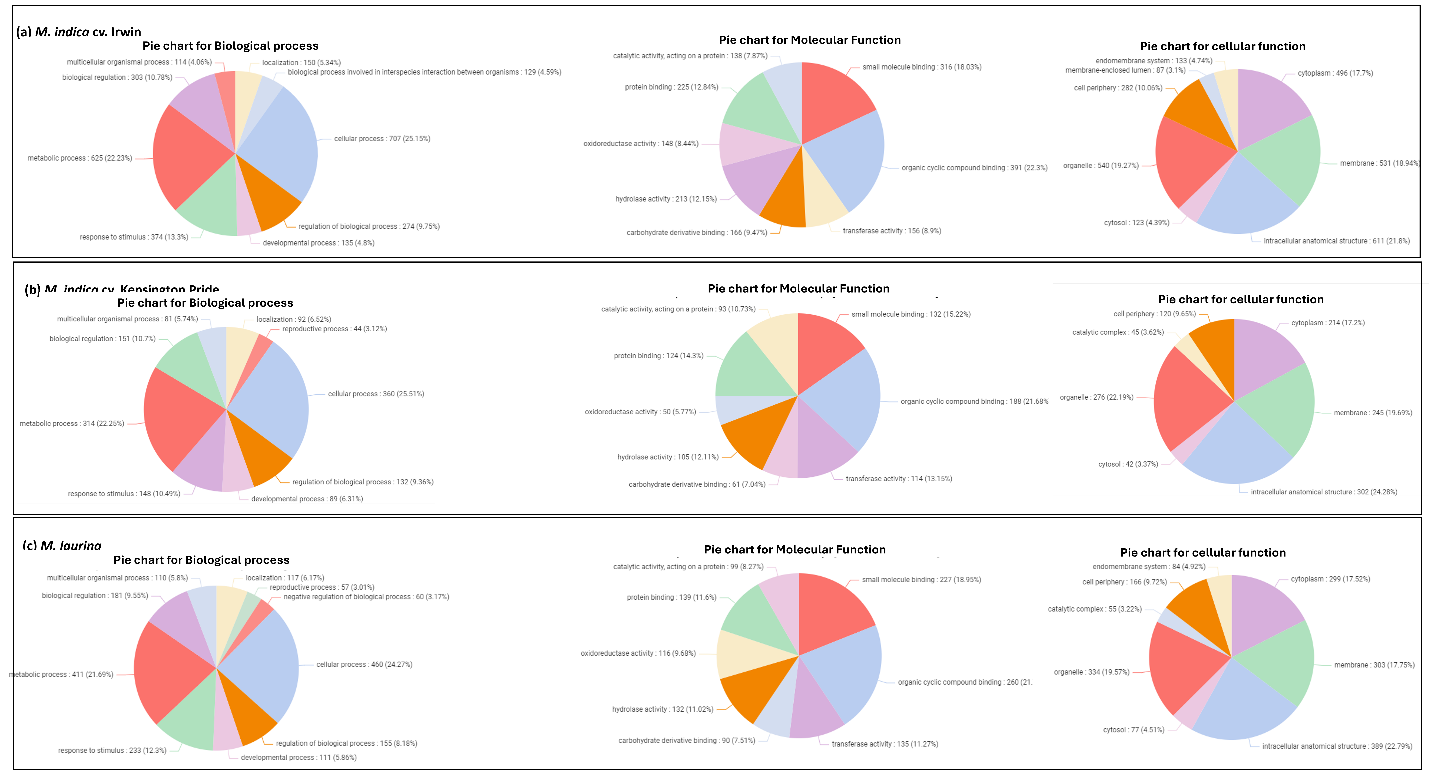

**Figure S7 Functions of unique genes in (a) Irwin, (b) Kensington Pride, and (c) *M. laurina* with respect to biological process, molecular function and cellular component.**

**Figure S8**
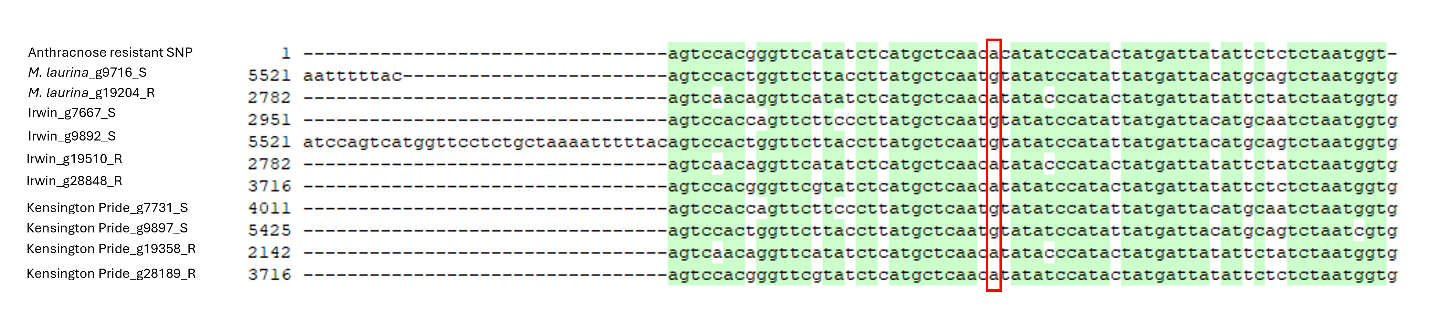
**Alignment of the region in β-1,3-glucanase 2 genes where the SNP for the anthracnose resistance is located.** SNP position is shown with red coloured box and species names and gene IDs are shown in the left side of the alignment.

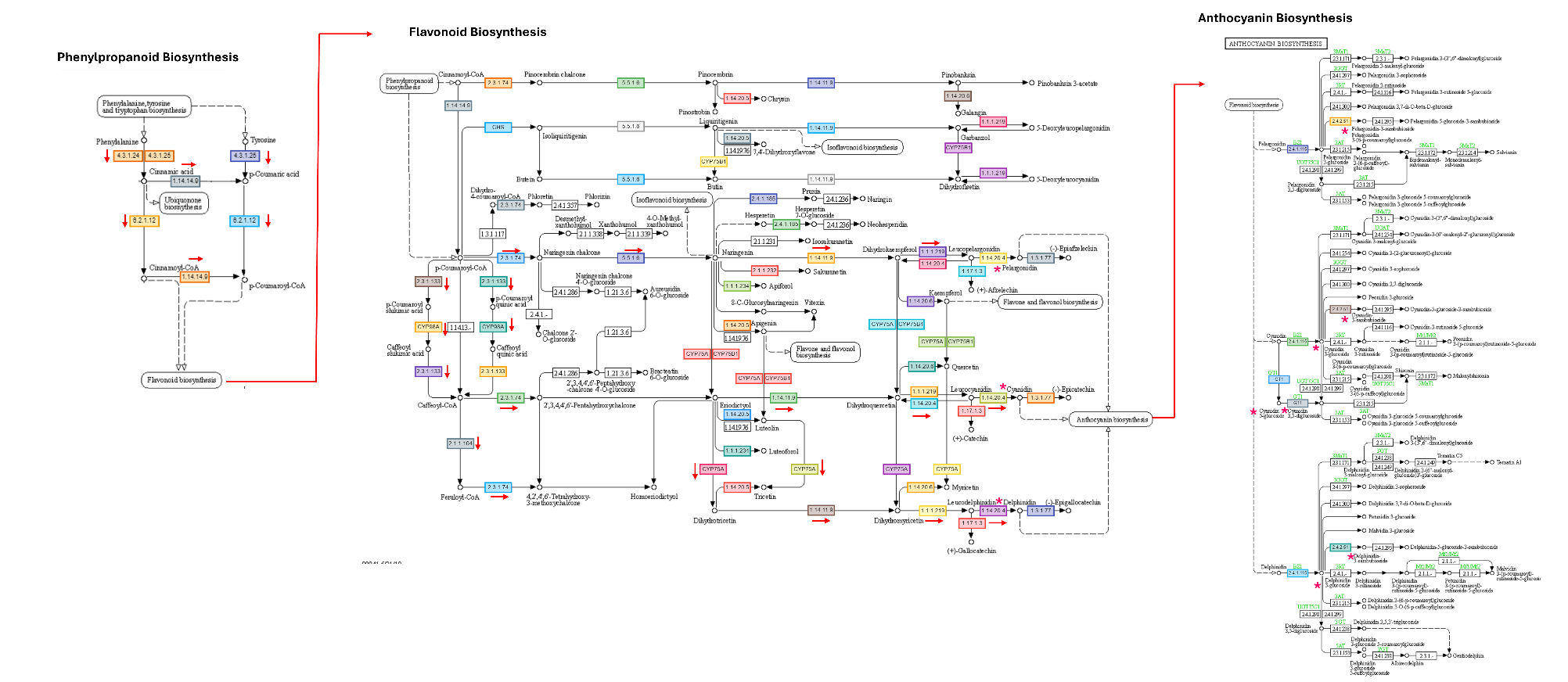

**Figure S9 Anthocyanin biosynthetic pathway of Irwin as a representative of all three genomes.** The main paths towards producing anthocyanins are shown with arrows and important anthocyanins produced are shown with red asterisks.
